# Mis-spliced transcripts generate *de novo* proteins in TDP-43-related ALS/FTD

**DOI:** 10.1101/2023.01.23.525149

**Authors:** Sahba Seddighi, Yue A. Qi, Anna-Leigh Brown, Oscar G. Wilkins, Colleen Bereda, Cedric Belair, Yongjie Zhang, Mercedes Prudencio, Matthew J Keuss, Aditya Khandeshi, Sarah Pickles, Sarah E. Hill, James Hawrot, Daniel M. Ramos, Hebao Yuan, Jessica Roberts, Erika Kelmer Sacramento, Syed I. Shah, Mike A. Nalls, Jenn Colon-Mercado, Joel F. Reyes, Veronica H. Ryan, Matthew P. Nelson, Casey Cook, Ziyi Li, Laurel Screven, Justin Y Kwan, Anantharaman Shantaraman, Lingyan Ping, Yuka Koike, Björn Oskarsson, Nathan Staff, Duc M. Duong, Aisha Ahmed, Maria Secrier, Jerneg Ule, Steven Jacobson, Jonathan Rohrer, Andrea Malaspina, Jonathan D. Glass, Alessandro Ori, Nicholas T. Seyfried, Manolis Maragkakis, Leonard Petrucelli, Pietro Fratta, Michael E. Ward

## Abstract

Functional loss of TDP-43, an RNA-binding protein genetically and pathologically linked to ALS and FTD, leads to inclusion of cryptic exons in hundreds of transcripts during disease. Cryptic exons can promote degradation of affected transcripts, deleteriously altering cellular function through loss-of-function mechanisms. However, the possibility of *de novo* protein synthesis from cryptic exon transcripts has not been explored. Here, we show that mRNA transcripts harboring cryptic exons generate *de novo* proteins both in TDP-43 deficient cellular models and in disease. Using coordinated transcriptomic and proteomic studies of TDP-43 depleted iPSC-derived neurons, we identified numerous peptides that mapped to cryptic exons. Cryptic exons identified in iPSC models were highly predictive of cryptic exons expressed in brains of patients with TDP-43 proteinopathy, including cryptic transcripts that generated *de novo* proteins. We discovered that inclusion of cryptic peptide sequences in proteins altered their interactions with other proteins, thereby likely altering their function. Finally, we showed that these *de novo* peptides were present in CSF from patients with ALS. The demonstration of cryptic exon translation suggests new mechanisms for ALS pathophysiology downstream of TDP-43 dysfunction and may provide a strategy for novel biomarker development.

**One Sentence Summary:** Loss of TDP-43 function results in the expression of *de novo* proteins from mis-spliced mRNA transcripts.

## INTRODUCTION

Cytoplasmic inclusions of the TAR DNA-binding protein 43 (TDP-43) occur in the brains and/or spinal cords of approximately 97% of amyotrophic lateral sclerosis (ALS), 45% of frontotemporal dementia (FTD), and 40% of Alzheimer’s disease *cases*(*1, 2*). Mutations in *TARDBP*, the gene encoding TDP-43, cause familial forms of FTD and ALS, further supporting a central role of TDP-43 in disease pathogenesis(*3*). TDP-43 mislocalization involves both its clearance from the nucleus and the formation of cytosolic aggregates(*1, 2, 4*). Nuclear clearance of TDP-43 can occur prior to symptom onset and precedes the formation of cytosolic aggregates(*5–7*), suggesting that loss of normal TDP-43 function is an early disease mechanism.

TDP-43 has two RNA recognition motifs and directly regulates RNA metabolism by acting as a potent splicing repressor. When TDP-43 splicing repression is lost, erroneous inclusion of intronic sequences called cryptic exons (CEs) occurs (*8*). Because CEs are non-conserved intronic sequences, they often introduce frameshifts, premature stop codons, or premature polyadenylation sequences. Such splicing errors can reduce expression of affected transcripts via nonsense-mediated decay (NMD) or other RNA degradation pathways(*9*). In turn, the protein levels of affected genes often decline in parallel. One such example is the well-characterized CE in stathmin-2 (*STMN2*), which causes loss of STMN2 mRNA and protein in the cortex and spinal cord of patients with ALS and FTLD-TDP(*10–12*). We and others recently discovered that TDP-43 also normally represses a cryptic exon in *UNC13A*, a critical synaptic *gene*(*13, 14*). ALS- and FTD-linked risk variants in *UNC13A* reduce affinity of TDP-43 for *UNC13A* transcripts, thereby promoting CE expression and *UNC13A* loss in neurons with TDP-43 insufficiency(*13*). These two examples clearly demonstrate that mis-splicing due to TDP-43 mislocalization can reduce expression of critical downstream genes, likely impacting neuronal biology during disease.

TDP-43 loss causes widespread cryptic exon expression and other related pathological splicing events that affect hundreds of transcripts (*8, 11, 12*). While certain genes exhibit reduced expression following CE inclusion, the possibility that *de novo* proteins could be synthesized from CE transcripts has never been addressed. Expression of *de novo* proteins downstream of TDP-43 loss of function could have several important ramifications. Translation of cryptic exons could alter the functions of proteins or induce toxic gain-of-functions with pathophysiological implications. Alternatively, *de novo* proteins could trigger an autoimmune response, akin to previous observations in autoimmune encephalitis and cancer(*15, 16*). Finally, discovery of *de novo* proteins caused by loss of TDP-43 function could enable the development of novel biomarkers of TDP-43 pathology and its functional loss.

Here, we systematically address whether loss of TDP-43 function leads to the expression of CE-encoded *de novo* polypeptides. Using a human iPSC-derived neuron model of TDP-43 loss-of-function, we developed a deep, high-quality neuronal CE atlas. We cross-referenced these CEs against a ribosome sequencing (Ribo-seq) dataset from TDP-43 knockdown (KD) neurons, discovering extensive ribosome-CE interactions that suggested their active translation. We then developed an unbiased proteogenomic pipeline, using coordinated short- and long-read transcriptomic and proteomics studies in iPSC neurons, which identified numerous peptides that mapped to cryptic exon coding frames. We identified these iPSC-predicted cryptic exons in FTD/ALS post-mortem cortex, validating the physiological relevance of our *in vitro* models. Through western blot and proteomic strategies, we confirmed the presence of *de novo* peptide expression in iPSC neurons and showed that they can alter the biology of proteins in which they are expressed. Finally, we developed a targeted proteomics assay to unequivocally identify *de novo* “cryptic” peptides in cerebrospinal fluid (CSF) from ALS patients. These observations of cryptic exon translation downstream of TDP-43 dysfunction both broaden our understanding of ALS pathophysiology and provide a new avenue for the development of molecular biomarkers for ALS and other neurodegenerative disorders.

## RESULTS

### Ribosomes bind to intronic regions of mis-spliced transcripts in TDP-43 deficient iNeurons

Ribo-seq provides a map of the position and density of ribosomes on individual mRNAs and a proxy for protein translation. If TDP-43 deficient neurons translate cryptic exons into *de novo* proteins (Fig. 1A), we reasoned that Ribo-seq may reveal ribosome-bound cryptic exons. Using short-read total RNA-seq in TDP-43 KD human iPSC-derived glutamatergic neurons (iNeurons, Fig. S1 A,B), we identified 340 abnormal splicing events across 233 genes (out of a total of 498 alternative splicing events, Fig. 1B, Fig. S1C, Table S1A). These abnormal splicing events included previously identified targets of TDP-43-related splicing repression, such as cryptic exons in *STMN2* and *UNC13A*(*11, 13*). The majority (70%) of cryptic splicing events were cassette exons, with the remainder comprising exon extension (12%), intron retention (9%), and exon skipping (9%) (Fig. S1C,D, Table S1A, File S2).

**Fig. 1.**
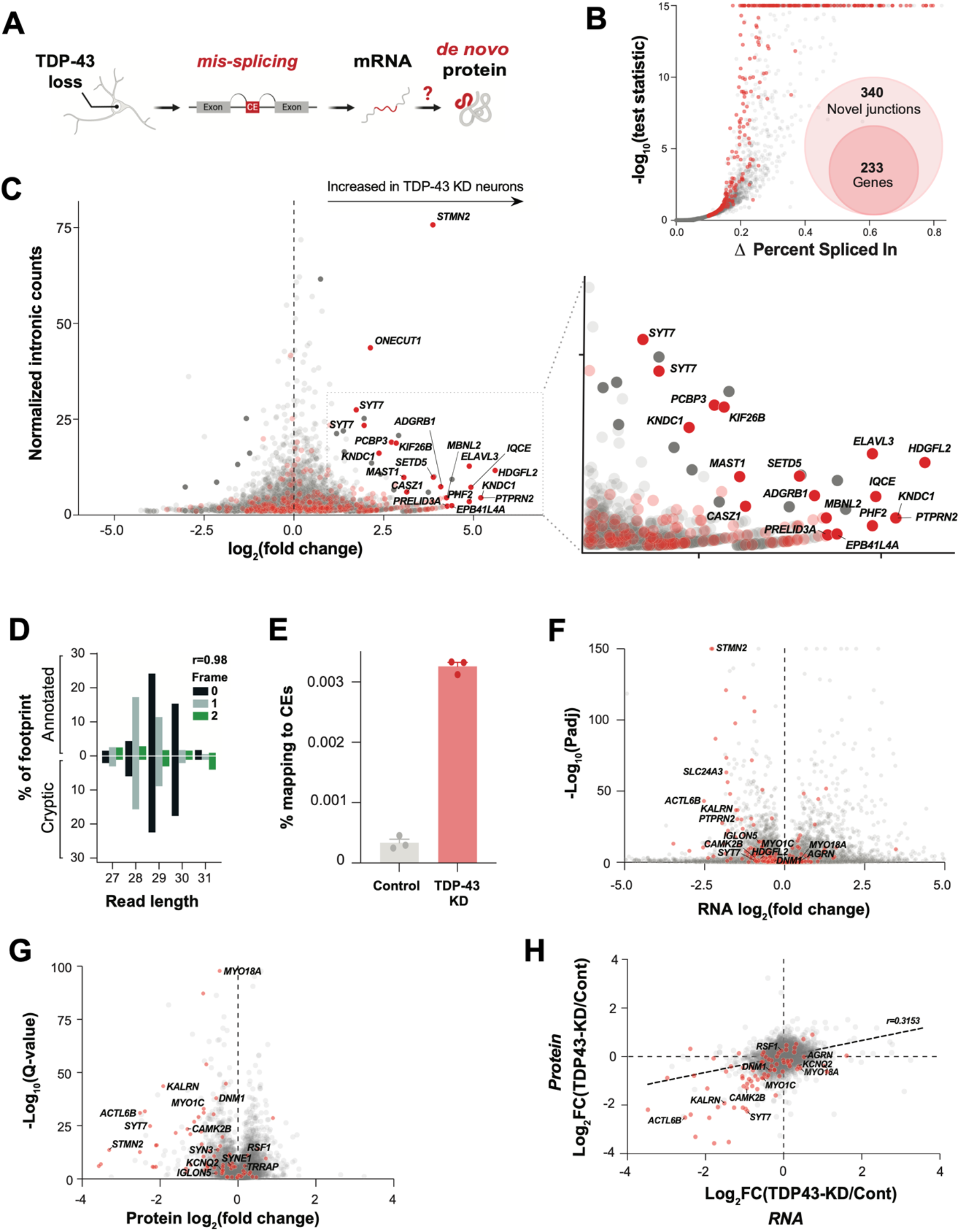
TDP-43 depletion causes ectopic ribosomal footprints that map to introns and alters the transcriptional and proteomic landscape in neurons. (**A**) Graphical overview of cryptic exon RNA and potential *de novo* protein production due to pathological mis-splicing in TDP-43 deficient neurons. (**B**) Differential splicing in TDP-43 KD versus control iNeurons. Cryptic exon genes are shown in red. (**C**) Differential ribosome footprint density of intronic mRNA in TDP-43 KD versus control iNeurons. Cryptic exon genes are shown in red. (**D**) Percentage abundance of each footprint type, defined by length and sub-codon position, for annotated coding sequences (positive Y-axis) and a subset of high-confidence cryptic exons which are predicted to be translated (negative Y-axis). The Pearson correlation between the values for annotated and cryptic transcripts is shown. Data is from one TDP-43 knockdown sample. (**E**) Percentage of footprints aligning to the subset of cryptic exons used in part C for three controls and three TDP-43 knockdown samples. (**F**) Differential transcript abundance in control and TDP-43 KD iNeurons, quantified using total short-read RNA-seq. Cryptic exon genes shown in red. (**G**) Differential protein abundance in control and TDP-43 KD iNeurons, quantified using mass spectrometry proteomics. Cryptic exon genes shown in red. (**H**) Quantification of mRNA (RNAseq) and protein (DIA proteomics) of control and TDP-43 KD iNeurons. Cryptic exons are shown in red.

We then queried iNeuron Ribo-seq data (*13*) for evidence of changes in ribosome binding within all intronic regions upon TDP-43 KD. We found a number of intronic sequences that were increased in TDP-43 KD Ribo-Seq (Table S1B). When we cross-referenced these sequences with CEs and other pathological splicing events identified in Figure 1B, we identified a substantial enrichment for CE-containing genes (Fig. 1C). Next, we selected CEs predicted to generate inframe peptides and investigated the periodicity of their ribosome footprints. Strikingly, these showed similar periodicity to that of all other constitutive exons, suggesting that the signal in these cases was genuinely derived from translating ribosomes (Fig. 1D). Overall, CEs were greatly enriched in TDP-43 KD Ribo-seq data compared to controls (Fig. 1E).

Using total RNA-seq and proteomics, we characterized the impact of neuronal TDP-43 loss on CE gene expression at the transcript and protein level (Fig. 1F-H, Fig. S1E-G, Table S1C,D). Consistent with prior observations that CEs can reduce mRNA stability(*10–14*), 37 percent of CE gene transcripts were reduced in TDP-43 KD neurons (Fig. 1F). We found evidence of additional instability of mis-spliced genes at the protein level, where 80 percent of proteins from CE genes displayed decreased expression (Fig. 1G,H). Collectively, these data show that while CEs often reduce expression of CE genes, certain CEs and other pathologically spliced transcripts might be translated into *de novo* proteins.

### TDP-43 deficient iNeurons express cryptic exons that generate *de novo* proteins

To identify potential CE-associated *de novo* proteins at a large-scale, we developed an unbiased proteogenomic pipeline (Fig. 2A). Using RNA-seq data from TDP-43 KD iNeurons, the pipeline returns in-frame amino acid sequences from mis-spliced junctions, henceforth referred to as cryptic peptides. We then added these *in silico* translated protein sequences of CEs to a standard human proteome reference (File S1), allowing us to search for trypsin-digested cryptic peptides in shotgun proteomic datasets from TDP-43 KD iNeurons. This pipeline identified 65 putative trypsin-digested cryptic peptides across 12 genes in TDP-43 KD iNeurons (Fig. 2B, Table S2A, Table S2B), consistent with the possibility that certain cryptic exons are translated into proteins.

**Fig. 2.**
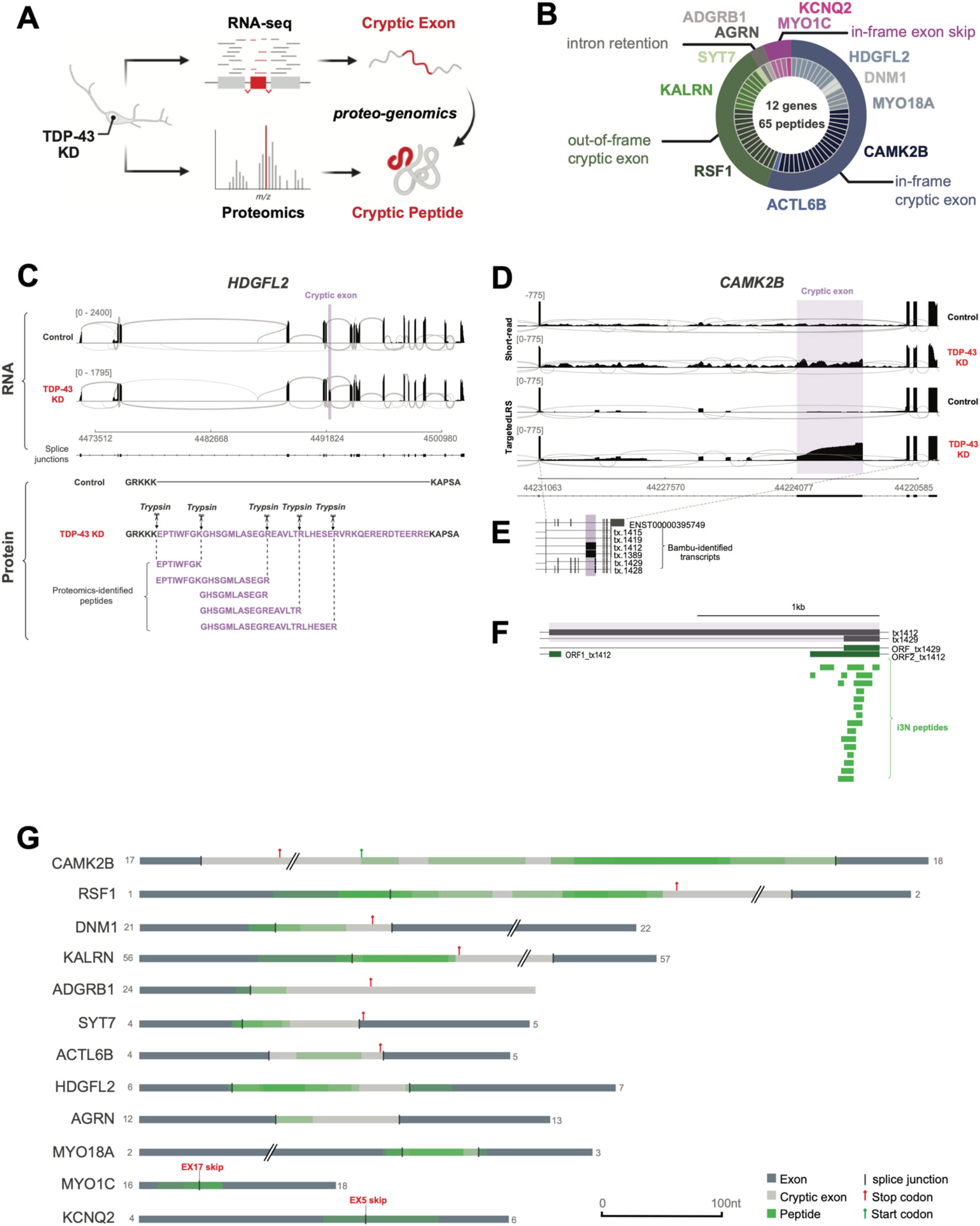
TDP-43 depletion causes formation of *de novo* cryptic peptides from mis-spliced transcripts in iPSC neurons. (**A**) Schematic of proteogenomic pipeline to identify *de novo* cryptic peptides caused by RNA mis-splicing in TDP-43 deficient neurons. (**B**) Proteogenomics of TDP-43 depleted iNeurons identified 65 putative trypsin-digested cryptic peptides across 12 genes. Outer circle of pie chart represents genes. Inner circle represents the number of putative trypsin-digested cryptic peptides that were identified for each gene. (**C**) Representative sashimi plots showing inclusion of an in-frame cryptic exon (purple) in HDGFL2 transcripts in TDP-43 deficient iNeurons. Amino acid sequence of putative translation of this cryptic exon shown below sashimi plot, with trypsin cleavage sites and proteogenomic-identified cryptic peptides annotated. (**D**) Representative sashimi plots comparing Illumina short-read RNA-seq versus Nanopore long-read sequencing of the *CAMK2B* transcript in control and TDP-43 KD iNeurons. Cryptic exon expressed in TDP-43 KD neurons is highlighted in purple. (**E**) Precise mapping of *CAMK2B* exon junctions and splice isoforms of transcripts in TDP-43 KD neurons using Nanopore long-read sequencing. (**F**) Tryptic-cryptic peptides (green) identified using the proteogenomic pipeline are mapped to an in-frame cryptic exon in *CAMK2B* identified using Nanopore long-read sequencing. (**G**) Graphical representation of proteogenomic-identified cryptic peptides mapped to transcripts from Nanopore long-read sequencing. Canonical upstream and downstream exons are in dark grey. Cryptic exons are in light grey. Cryptic peptide locations are overlaid in green.

More than half of the identified cryptic peptides were predicted to be in frame when mapped to short-read RNA-seq of pathologically spliced transcripts (Fig. 2B). One such example is a 46-amino-acid, in-frame cryptic peptide in HDGFL2 that mapped to a CE in intron 6 (Fig. 2C, S2A). We detected five distinct trypsin-cleaved peptides for HDGFL2 in TDP-43 KD iNeurons (Fig. 2C). Multiple out-of-frame cryptic peptides were also detected and mapped to cryptic exons in RSF1, KALRN, and SYT7 (Table S2A). Based on the frame of these out-of-frame cryptic peptides, the parent transcripts were predicted to harbor a premature stop codon in the downstream exon. A smaller fraction of cryptic peptides either mapped to retained introns or were due to exon skipping (Fig. 2B).

Short-read RNA-seq relies on informatic approaches to stitch together individual reads into full-length transcripts. Errors in predicted full-length sequences can arise when multiple transcript isoforms are co-expressed, particularly in settings of pathological splicing. To unequivocably determine the frame of translation for each cryptic peptide, we performed Nanopore long-read sequencing of control and TDP-43 KD iNeurons (Fig 2D-G, S2B-D, Table S2C). Mapping all putative cryptic peptides against long-read data from which we could extract the frame confirmed six genes with in-frame cryptic exons in TDP-43 KD neurons (Fig 2G). An example of an in-frame cryptic peptide from *CAMK2B* is shown in Figure 2D-F, demonstrating the improved fidelity of long-read sequencing over short-read sequencing in mapping cryptic peptides to complex, pathologically spliced transcripts. Taken together, these data provide additional evidence that certain cryptic exons and other pathologically spliced transcripts likely generate *de novo* proteins in TDP-43 deficient neurons.

### TDP-43 deficient iPSC neurons accurately predict mis-splicing in ALS/FTD patient brains

Previously, TDP-43 deficient human iPSC neuronal models successfully predicted individual cryptic exons that were later validated in FTD/ALS patients(*11–13*). Most of the putative cryptic peptides that we observed mapped to cryptic exons that have not been previously described. We therefore tested the fidelity of our iPSC neuron model in global prediction of FTD/ALS associated cryptic exons, using three separate approaches. First, we cross-referenced cryptic exons found in TDP-43 KD iNeurons against a published RNAseq dataset of FACS-sorted neuronal nuclei from FTD/ALS cortex (*17*). Of the 340 pathological splice junctions that we identified in TDP-43 KD iNeurons, 230 were detected in FACS-sorted post-mortem neuronal nuclei. Of these, 122 junctions were enriched in nuclei lacking TDP-43 (Fig. 3A, Table S3A,B). Using dimensionality reduction analysis of iNeuron-predicted cryptic exons, we found that TDP-43 positive and negative nuclei clearly separated based on the first principal component, accounting for 47.01% of the variability in the dataset (Fig. 3B). These data indicate that numerous cryptic splicing events observed in TDP-43 deficient iPSC-derived neurons also occur in TDP-43 deficient ALS/FTD patient neurons.

**Fig. 3.**
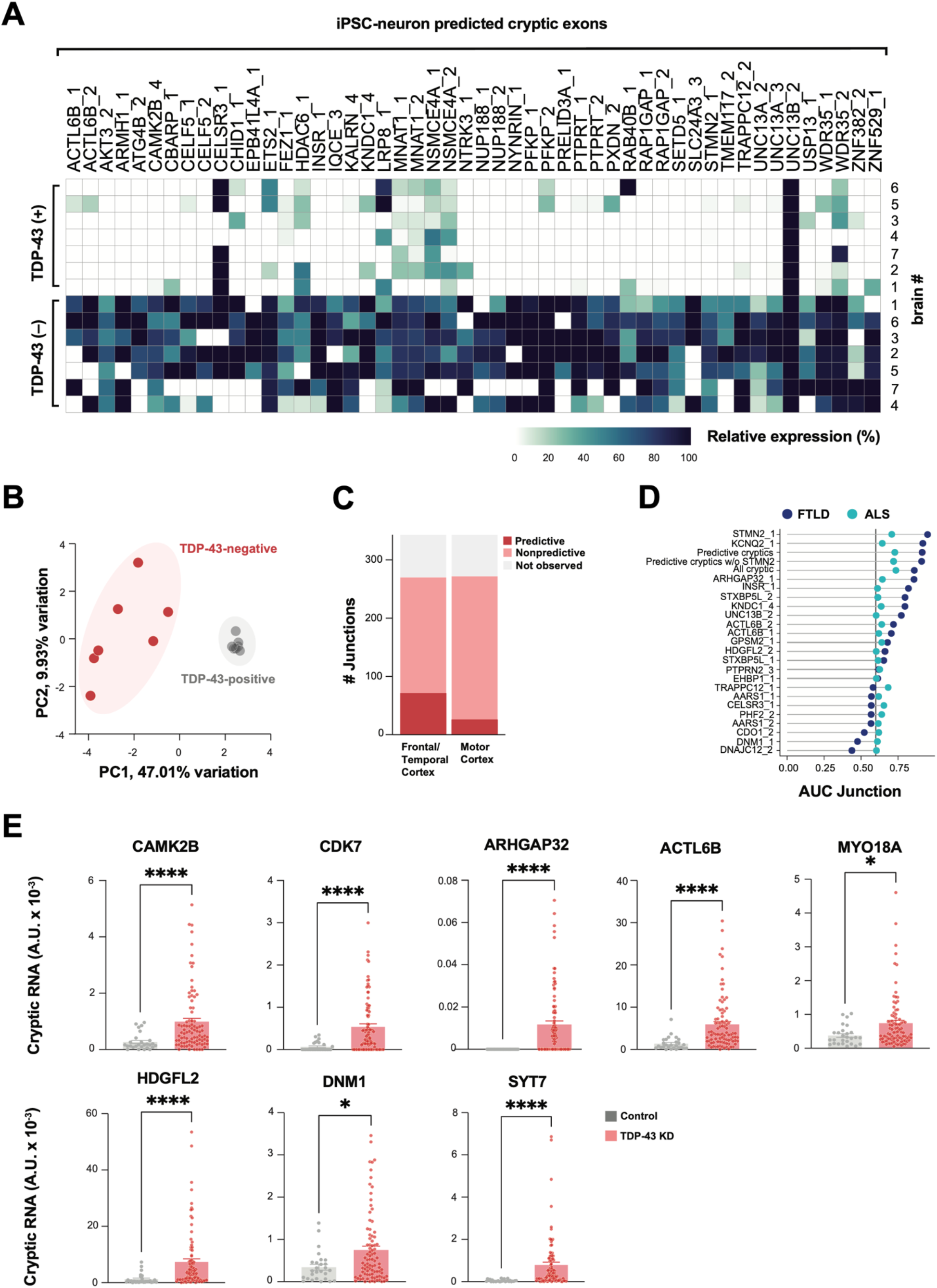
TDP-43 cryptic exons identified in iPSC neurons classifies TDP-43 pathology in ALS/FTD CNS tissue. (**A**) Heatmap of percent spliced in (PSI) of iNeuron-predicted cryptic exons in FACs-sorted TDP-43 positive and negative neuronal nuclei from postmortem FTD/ALS cortex (*17*). We defined enriched junctions as those with PSI in TDP-43 negative nuclei greater than twice the PSI in TDP-43 positive (mean TDP-43 negative PSI > 0.10). The top 50 most expressed cryptic splice junctions in TDP43-negative nuclei, compared to TDP-43 positive nuclei, are shown, and cases were organized by unsupervised hierarchical clustering based on cryptic exon PSI. (**B**) Principal component analysis (PCA) on PSI of 230 iNeuron-predicted TDP-43 cryptic exons in TDP-43 positive and negative neuronal nuclei from FTD/ALS cortex. (**C**) Bar plot showing the number of iNeuron-predicted cryptic exons detected in bulk postmortem RNA-seq from a NYGC ALS/FTD. Cryptic exons are classified by whether they were observed in postmortem tissue, detected and non-predictive, or predictive between TDP-43 pathology from non-TDP-43 pathology, as shown in D, with an area under the curve (AUC) >= 0.6. (**D**) AUC of iNeuron-predicted cryptic exons for classification of TDP-43 pathology in FTLD frontal/temporal cortex and ALS motor cortex postmortem tissue. Meta-expression score of all cryptic exons, of only predictive exons (AUC >= 0.6), and of predictive exons excluding *STMN2* expression were also tested. (**E**) QRTPCR-based validation of eight cryptic exons in an independent cohort of FTLD-TDP frontal cortex (n=89 FTLD-TDP cases, n=27 non-neurological disease controls). Data are presented as mean +/- SEM. *P*-values shown resulting from Mann-Whitney test: \**p*≤0.05, \*\**p*≤0.005, *** *p*≤0.0005; \*\*\*\**p*≤0.0001.

Second, we tested if cryptic splicing predicted by iNeurons could identify cases with TDP-43 pathology in post-mortem cortex. We analyzed bulk RNAseq datasets from the New York Genome Center (NYGC), comprising 168 frontal and temporal cortex samples from 82 non-neurological disease controls, 20 FTLD-non-TDP, and 66 FTLD-TDP, and 304 motor cortex samples from 49 non-neurological disease controls, 11 ALS non-TDP, and 244 ALS-TDP. Despite the anticipated low expression and degradation of many cryptic exons by nonsense-mediated decay (NMD), and the limits of bulk RNA-seq in which only a small proportion of cells have TDP-43 pathology, we detected 298 of the 340 pathologically spliced junctions with at least one spliced read across all the samples (Fig. 3C). Next, we tested if CE expression levels could predict cases with TDP-43 proteinopathy. Numerous CE junctions identified in TDP-43 deficient iNeurons had positive prediction power in FTLD (69) and ALS (25) (Fig. 3C, Table S3A,C,D). We then generated meta-scores by combining the expression levels of individual or multiple cryptic junctions. We created three different meta-scores: One using the expression of all cryptic events, one using only those junctions with a high predictive power (AUC >= 0.6), and one with all predictive cryptic junctions excluding the *STMN2* junction. In the FTLD and ALS samples, *STMN2, KCNQ2*, and all predictive cryptic junctions had the highest power to distinguish cases from controls (Fig. 3D, Table S3A,C,D).

Third, we used quantitative PCR with reverse transcription (RT-qPCR) to assess CE expression in an independent cohort of 89 FTD/ALS postmortem cortex versus 27 healthy controls, focusing on eight transcripts associated with potential cryptic peptide expression (Table S3E). We observed substantially higher expression of each of these cryptic exons in patients compared to controls (Fig. 3E, Fig. S3A). These three independent approaches show that dysregulated splicing across numerous transcripts can be accurately predicted by TDP-43 deficient iNeurons, justifying physiological relevance of their potential translation into *de novo* proteins.

### Expression of cryptic exons alters protein interactomes

To validate the existence of a *de novo* protein sequence identified by proteogenomics, we developed an antibody against an *in-frame* cryptic peptide in MYO18A. This peptide was predicted to be both highly polar and on the protein surface based on an AlphaFold structural model (Fig. 4A, Fig. S4A). Using this antibody in a western blot assay, we detected a protein of the anticipated molecular weight of MYO18A expressing a cryptic exon (MYO18A-CE) in TDP-43 KD, but not control, iNeurons (Fig. 4B,C). We next assayed HDGFL2, a protein modestly reduced in expression in TDP-43 deficient neurons (Table S1D) and predicted to express an in-frame cryptic peptide (Fig. 2C). A commercially available polyclonal antibody against canonical HDGFL2 detected a single band on western blot at the predicted molecular weight in control neurons (Fig. 4D). However, this antibody also detected a second, higher-molecular protein in TDP-43 KD iNeurons, matching the predicted molecular weight of HDGFL2 expressing a cryptic exon (HDGFL2-CE) (Fig. 4D,E). These data validate our predictions that some cryptic exons are translated into *de novo* proteins expressed highly enough to be detected by standard immunoassays.

**Fig. 4.**
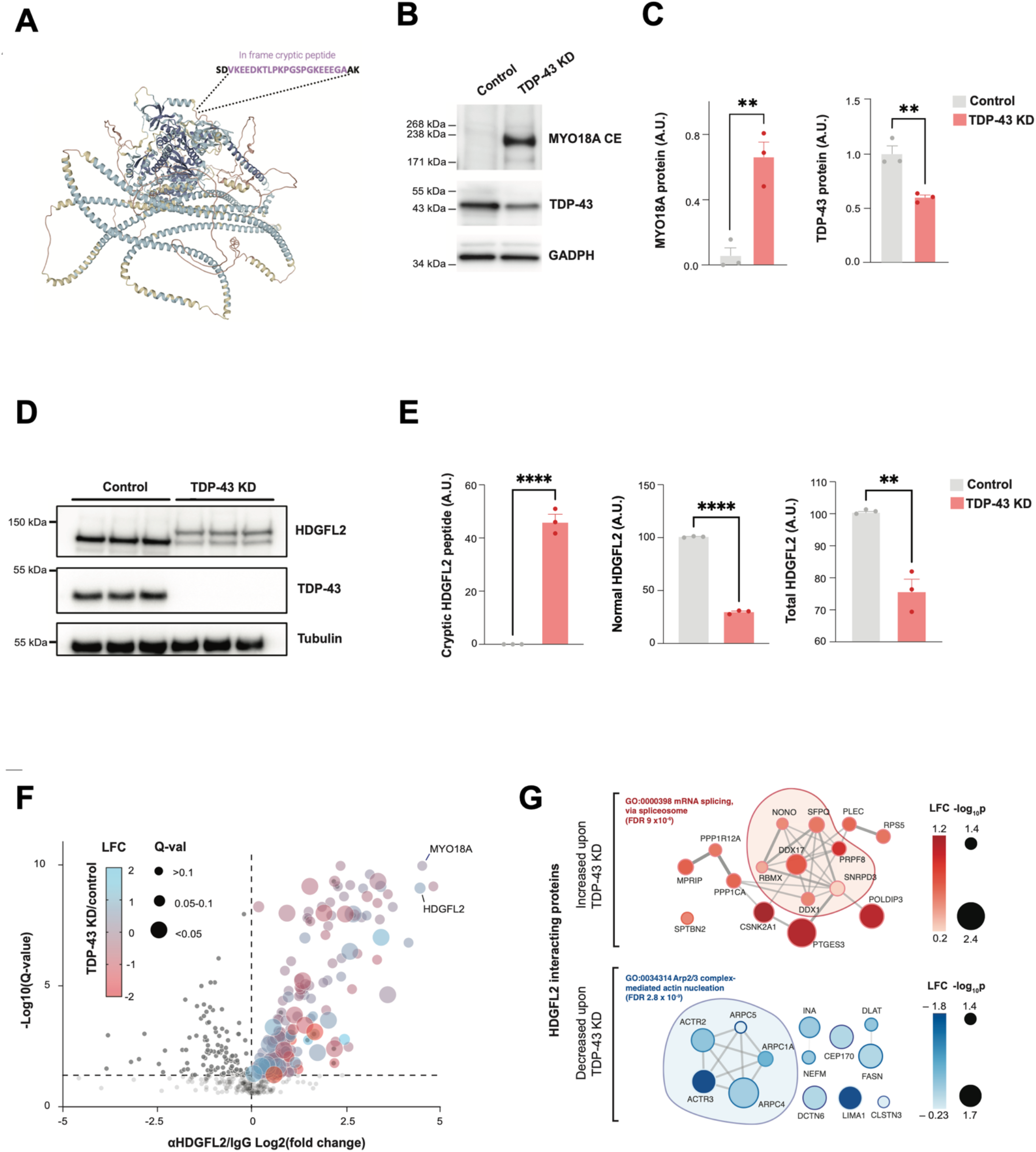
Cryptic peptides in TDP-43 deficient iPSC neurons can be orthogonally validated and alter the physical interactomes of the proteins in which they are expressed. (**A**) Predicted structure of MYO18A (AlphaFold), annotated with the predicted insertion site of a polar, in-frame cryptic exon on the protein surface, which is induced by TDP-43 depletion. A polyclonal antibody was raised against this *MYO18A* cryptic exon. (**B**) Antibody-based detection of a MYO18A cryptic peptide in TDP-43 depleted iNeurons. Representative western blot showing a band of the expected molecular weight of MYO18A-CE specifically in TDP-43 depleted neurons. (**C**) Quantification of MYO18A-CE and TDP-43 expression levels (n=3, 2-sample t-test, **p<0.01). (**D**) Representative western-blot detection of a higher molecular weight species of HDGFL2 protein in TDP-43 knockdown iPSC neurons, using an antibody against total HDGFL2. Higher molecular weight species of HDGFL2 is the approximate size of the canonical protein plus the proteogenomic-predicted in-frame cryptic exon. (**E**) Quantification of HDGFL2-CE, HDGFL2, and HDGFL2+HDGFL2-CE (n=3, 2-sample t-test, **p<0.01, ****p<0.0001). (**F**) Affinity purification mass spectrometry analysis of HDGFL2 protein-protein interactions in control and TDP-43 depleted iPSC neurons. Volcano plot of co-immunoprecipitated proteins using anti-HDGFL2 antibody versus control IgG is shown. Dot color reflects LFC in TDP-43 KD versus control neurons, and dot size reflects the adjusted *p*-value. (**G**) STRING diagram of proteins whose interactions with HDGFL2 are significantly altered by TDP-43 knockdown. Dot color reflects LFC in TDP-43 KD versus control neurons.

We reasoned that inclusion of a 46-amino-acid cryptic peptide in HDGFL2, a protein thought to regulate chromatin and DNA repair in non-neuronal cells (*18*), might alter its interacting partners and thus its biology. We performed affinity purification mass spectrometry (APMS) of HDGFL2 in control and TDP-43 KD iNeurons, using an anti-HDGFL2 antibody to immunoprecipitate HDGFL2 and its associated proteins. We identified 178 HDGFL2-interacting proteins that were significantly enriched in anti-HDGFL2 compared to control IgG pulldown conditions (Fig. 4F, Table S4A,B). GO-term analysis of HDGFL2 interacting proteins in neurons revealed strong enrichments in ribosome, spliceosome, actin, and neurodegeneration-related pathways (Fig. S4B,C). Surprisingly, MYO18A was a top interacting protein of HDGFL2, clustering with other actin-regulating HDGFL2 interactors (Fig. S4D). We then analyzed how TDP-43 loss – which causes expression of HDGFL-CE – altered the HDGFL2 interactome. Sixteen proteins increased their relative interactions with HDGFL2 upon TDP-43 loss, a large fraction of which regulate mRNA splicing, chromatin remodeling, and DNA repair (Fig. 4G). In contrast, thirteen proteins decreased their relative interactions with HDGFL2 upon TDP-43 loss. Most of these proteins play roles in cytoskeleton organization, with five comprising core regulators of Arp2/3 actin nucleation. These observations suggest that inclusion of cryptic exon sequences in a translated protein can alter its protein-protein interactome downstream of TDP-43 loss of function.

### Scalable validation of cryptic peptides by targeted proteomics

Because antibody development against novel peptides is a time and resource-intensive endeavor, we developed a targeted mass spectrometry-based approach to validate and measure additional cryptic peptides predicted by proteogenomics. By co-injecting endogenous cryptic peptides and a panel of their stable isotope (SIL) heavy peptide standards coupled with parallel reaction monitoring (PRM) mass spectrometry proteomics, one can specifically quantify dozens of peptides in a single sample run (Fig. 5A). We designed PRM assays for the 65 cryptic peptides predicted by our proteogenomic pipeline (Table S5A). Of these, we successfully quantified twelve endogenous trypsin-digested cryptic peptides across four genes in TDP-43 KD iNeurons (Fig. 5B-D, S5A) that were precisely coeluted with their SIL counterparts (Fig. 5B, S5A), with almost identical fragmentation spectra (Fig. 5C, S5A). Consistent with cryptic exon mRNA expression, we found that cryptic peptides from all four genes were highly increased in TDP-43 KD iNeurons, with nearly no expression in control neurons (Fig. 5E, Table S5B). These experiments provide further evidence that multiple genes express cryptic peptides in the setting of functional TDP-43 loss, and demonstrate that proteomics can scalably detect and measure the expression of cryptic peptides in complex biological samples.

**Fig. 5.**
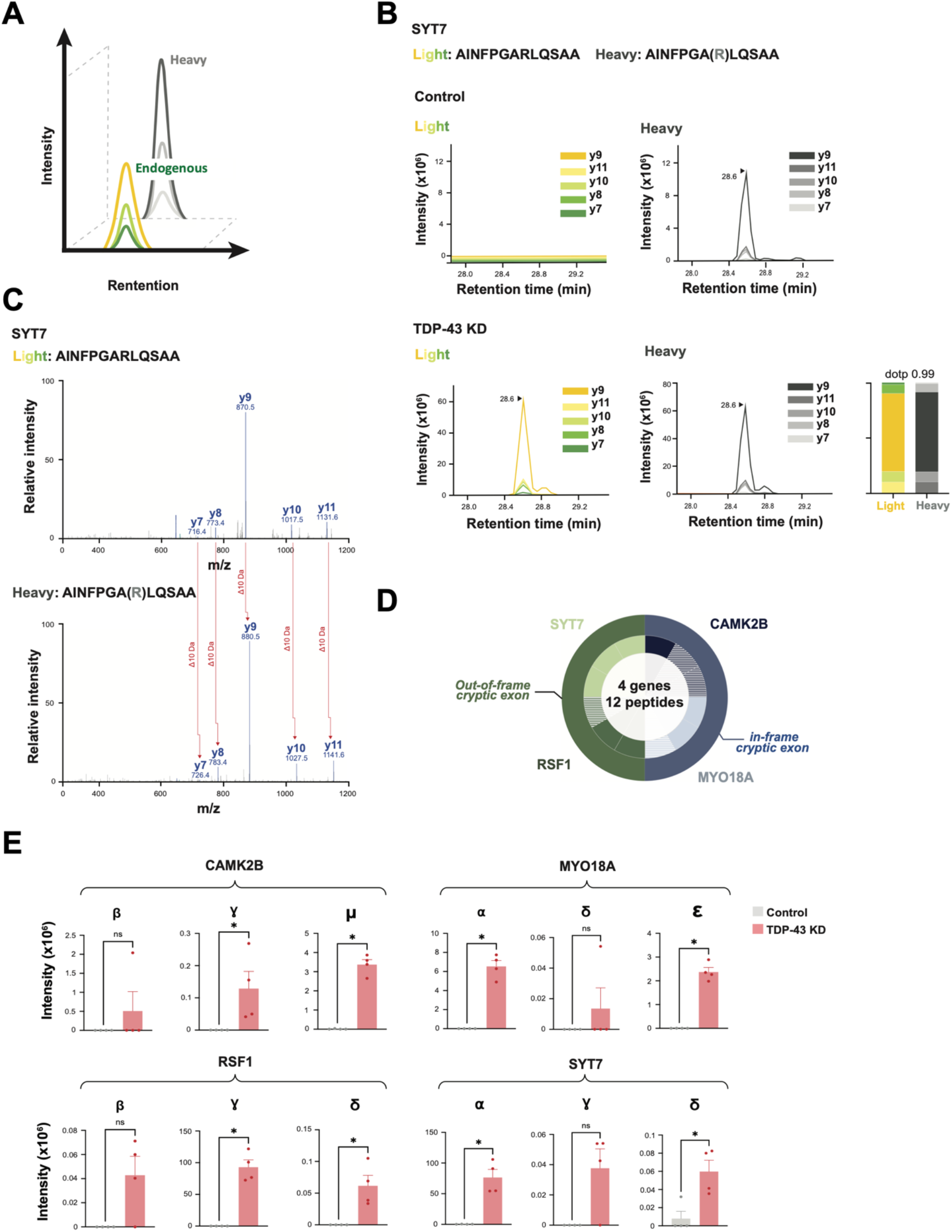
Scalable cryptic peptide validation in TDP-43 deficient neurons by targeted proteomics. (**A**) Schematic of parallel reaction monitoring targeted mass spectrometry (PRM-MS). Co-elution of stable heavy isotope-labeled peptides (SIL peptides) allows for ultrasensitive measurement of corresponding endogenous peptides. (**B**) PRM-MS assay using a synthetic SIL peptide internal standard identifies a cryptic peptide in SYT7 in TDP-43 KD, but not control, iNeurons. Spectral plot of heavy standards and light (endogenous) y-ions from an SYT7 cryptic peptide are shown, with accompanying dot plots. (**C**) Corresponding mass spectra of endogenous and heavy peptide standards of the SYT7 cryptic peptide in TDP-43 depleted iPSC neurons. (**D**) Detection of 12 trypsin-digested cryptic peptides across 4 genes using singleshot PRM assays in TDP-43 KD iPSC neuron lysates. Outer circle represents the gene, and inner circle represents the number of cryptic peptides detected by PRM per gene. Hatched color signifies successful detection of 1-2 y-ions; solid color signifies detection of 3 or more y-ions. (**E**) Quantification of cryptic peptide expression in control and TDP-43 KD iPSC neurons using PRM assays. n= 4 replicates per sample. Two-sample t-test. *p<0.05, **p<0.01, ***p<0.001, ****p<0.001

### Identification of cryptic peptides in human CSF

We next asked if patients with FTD-ALS spectrum disorders associated with TDP-43 mislocalization express cryptic peptides. We compared our iNeuron cryptic exon dataset and a shotgun proteomics dataset from ALS CSF, with a goal of identifying genes that are both highly expressed at the protein level in ALS and could express cryptic exons in settings of TDP-43 loss. We discovered that forty-seven cryptic exon genes are present at the total protein level in CSF, including several that express cryptic peptides in TDP-43 deficient iNeurons (Fig. 6A,B, Table S6A). We searched two additional ALS CSF proteomics datasets from independent patient cohorts, thereby identifying four additional cryptic exon-containing genes with parent proteins in human CSF (Fig. 6B, Fig. S6A,B, Table S6B-E).

**Fig. 6.**
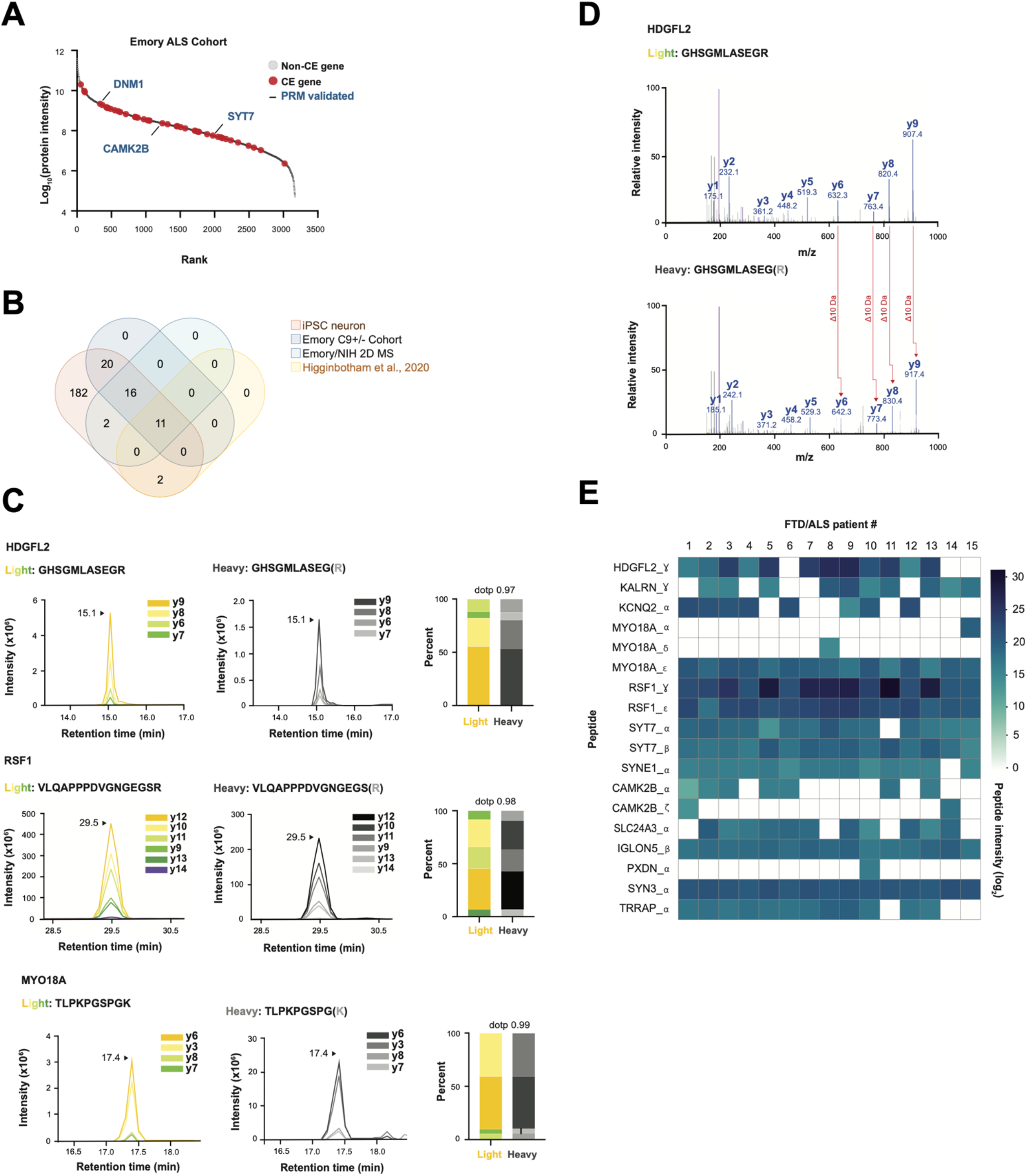
Cryptic peptides are present in human CSF. (**A**) Rank plot of proteins detected in CSF from patients with ALS. 47 cryptic exon genes that were predicted in TDP-43 KD iPSC neurons are shown in red. PRM-validated cryptic peptides expressing genes (from iNeuron studies) are annotated with blue text. (**B**) Venn Diagram of three CSF proteomics datasets versus cryptic exons predicted in TDP-43 KD iPSC neurons. (**C**) Representative spectral plots of three heavy (standard) and light (endogenous) cryptic peptides detected in ALS patient CSF. (**D**) MS/MS spectrum of a cryptic peptide in HDGFL2 that corresponds to the reference peptide (top) and endogenous peptide (bottom) detected in ALS patient CSF. (**E**) Heatmap of AUC intensities of 18 cryptic peptides from 13 different proteins in CSF from 15 ALS patients.

We augmented our previous list of SIL peptide standards with these additional candidates and used DIA proteomics to co-measure 65 endogenous and heavy standard standards in FTD/ALS patient CSF in a single MS run (Fig. S6C, Table S5A). We successfully detected eighteen peptides across thirteen genes that mapped to cryptic exons in FTD/ALS patient CSF (Fig. 6C-E, S7A, Table S6F,G). These endogenous tryptic-cryptic peptides co-eluted with their heavy peptide standards (Fig. 6C, S7A) and had matching MS/MS spectra (Fig. 6D). We monitored the expression of these peptides in CSF from fifteen patients with FTD/ALS. Most peptides were observed in multiple patients, with ten peptides across eight genes detectable in >80% of patients (Fig. 6E, Table S6F,G). These data indicate that, like TDP-43 KD iNeurons, patients with FTD/ALS spectrum disorders associated with functional TDP-43 depletion express and translate cryptic exons.

## DISCUSSION

TDP-43 loss of function causes widespread cryptic splicing that can reduce the expression of mRNA transcripts. Here, we provide the first demonstration that such mis-splicing can also generate *de novo* proteins from pathological transcripts. Our studies indicate that inclusion of cryptic exons in proteins can, in certain instances, alter the interacting partners of protein products. We show that human iPSC neurons accurately model hundreds of cryptic exons found in FTD/ALS brains, enabling prediction of cryptic peptides and development of antibody and mass spectrometry-based methods for their detection. Finally, we demonstrate the presence of TDP-43 related *de novo* peptides in CSF from ALS patients.

These discoveries are relevant to both FTD/ALS pathophysiology and diagnostics. In addition to the well-described loss-of-function mechanisms downstream of cryptic splicing, new possibilities of altered protein function should now be considered. For example, we found that TDP-43 loss increased HDGFL2 interaction with splicing regulators, while decreasing its interaction with other protein networks, such as those that regulate actin dynamics. This suggests both gain and loss of function of HDGFL2 occurs as a direct consequence of cryptic exon expression. Of note, we additionally found that HDGFL2 interacts with two other targets of TDP-43-related splicing regulation: MYO18A, a CE-harboring member of the actin regulatory network, and POLDIP3, which exhibits exon-skipping in settings of TDP-43 loss. Together, these observations hint at the possibility that TDP-43 loss of function may alter the biology of functionally interrelated proteins. Because transient mislocalization of TDP-43 occurs in settings of cell stress (e.g., following axotomy or oxidative stress), these findings imply that cryptic exon expression downstream of nuclear TDP-43 loss could play a regulatory role in settings outside of disease pathophysiology (*19–21*).

Our data suggest that only a small fraction of CE-harboring transcripts produce stable polypeptides, while most undergo degradation at the RNA and/or protein level. Of 154 CE-harboring genes, only 27 contained in-frame insertions, which are less likely to trigger RNA degradation pathways, such as nonsense-mediated decay. Additional quality control pathways may regulate the degradation of misfolded proteins translated from CEs. Despite high ribosome occupancy of *STMN2* intronic sequences, no peptides from *STMN2* were identified from our proteomic studies, suggesting that any proteins produced from the *STMN2* CE are highly unstable.

Our findings also open intriguing possibilities for a potential role of cryptic peptides in autoimmune dysregulation in neurodegenerative disorders. It is conceivable that cryptic peptides could be presented by MHCs for recognition by cytotoxic T-cells. Indeed, infiltration of cytotoxic CD8+ T cells has been previously reported in the brain and spinal cord of ALS patients(*22–24*). The possibility that cryptic peptides could elicit expression of autoantibodies is also possible. Neo-antigens are now well-described for other neurodegenerative disorders, including Lewy body dementia and Parkinson’s disease, and suggest the possibility of searching for T-cell or antibody-based responses against such antigens(*25, 26*). Retrospective analysis of stored serum samples recently led to the seminal discovery of EBV as the viral etiology of *MS*(*27, 28*). Similar approaches may be fruitful for ALS.

To accurately map putative cryptic peptides to mRNA transcripts, we generated the first-ever transcriptome-wide database of long-read sequencing in TDP-43 KD cells. While informatic methods to analyze long-read data for splicing abnormalities are still in development, we anticipate that these datasets will be of immediate use for the community. Peptide libraries can be generated from our proteomic datasets and used to probe for autoreactive B or T-cell populations and antibodies in patient biospecimens. The expanded identification of cryptic exon-harboring transcripts within postmortem patient brains may prove useful in understanding the pathophysiology of ALS/FTD and could be leveraged in the development of mRNA-based diagnostics. We provide validated amino acid sequences for cryptic peptides, which could serve as the basis for designing high-sensitivity antibody- or MS-based assays (eg, Simoa, SRM, IP-MS).

Currently, there are no biomarkers that enable identification of living patients with TDP-43 pathology, nor any means to monitor responses to TDP-43 directed therapies that are in development for FTD-ALS spectrum disorders. Approximately 20%-50% of AD patients – and 75% of patients with severe AD – also have TDP-43 co-pathology and may represent an additional subpopulation that could benefit from drugs targeting the TDP-43 *axis*(*29–31*). Identification of imaging-based biomarkers to detect TDP-43 or its pathological forms has proven challenging. Antibody-based strategies to monitor the levels and biochemical properties (eg, phosphorylation, solubility) of TDP-43 have been attempted; however, their interpretability and theragnostic use is challenged by the lack of a connection to the functional state of TDP-43 protein. A critical unmet need to develop molecular biomarkers of TDP-43 pathology therefore exists, both to guide disease stratification and to enable better deployment of TDP-43 directed therapies.

Since mis-splicing is a direct consequence of TDP-43 dysfunction, we reason that cryptic peptide biomarkers could be ideal for monitoring therapeutic responses. Targeted mass spectrometry-based approaches could be ideal for developing ultra-sensitive bioassays for clinical application. An example of this is selected reaction monitoring (SRM), wherein synthetic stable isotope-labeled peptides are used as internal standards to enable quantification of trace *analytes*(*32*). Alternate strategies may leverage high-sensitivity antibodies or aptamers against native proteins expressing cryptic peptides. Such tools could subsequently enable single molecular array (SIMOA) assays that can measure analytes at the single molecule *level*(*33*). Novel methods to enrich for brain-derived exosomes in plasma and pair with single molecule detection strategies (eg, SMAC) are being developed and open exciting possibilities for non-invasive disease profiling in *blood*(*34*). Ultimately, a multi-analyte panel for cryptic peptides may be ideal. This is consistent with our findings of improved disease predictability in postmortem brains upon consideration of multiple CEs over any single mis-spliced junction (except for *STMN2* CE, which cannot yet be detected at the protein level).

Our study has limitations. We did not evaluate whether the levels of cryptic peptides are elevated in cases compared to healthy controls. Our MS assays were designed to enable specific detection of target peptides; however, in their current form, they are only semi-quantitative and subject to technical artifacts when analyzing absolute levels of peptides in biofluid samples with high dynamic ranges and low peptide abundances. Our assays also require depletion of the top 14 contaminants in CSF, which introduces additional technical variability to sample preparation. SRM assays are the gold-standard for absolute protein quantification and enable target peptides to be measured without immunodepletion, but the development and optimization of such assays is highly laborious and specialized. The development of cryptic peptide biomarkers is further challenged by the current lack of a definitive antemortem TDP-43 assay. Perceived “healthy” controls could harbor early TDP-43 ‘opathies. Approximately 20-50% of all AD cases exhibit concomitant TDP-43 ‘opathy before overt symptom-onset(*29, 30*). Additionally, there are no clearly defined antemortem pathological markers of limbic predominant age-related TDP-43 encephalopathy (LATE). These challenges underscore a need for careful selection of control populations. Genetic markers, such as *C9orf72* repeat expansion, can be used to predict the presence of TDP-43 aggregates or suggest their absence in the setting of rare genetic markers, such as *FUS* and *SOD1* mutations. As a pure tauopathy, progressive supranuclear palsy may represent a superior control population to age-matched “healthy” individuals. With the current datasets at-hand, our hope is that multiple efforts to develop either mass spectrometry or antibody-based methods for absolute quantification can ensue in parallel.

In summary, we provide the first demonstration of *de novo* cryptic peptides occurring downstream of TDP-43 dysfunction in cellular models and patient biofluids. These findings should guide new discoveries on the role of cryptic peptides in TDP-43 ‘opathies and provide a framework for the development of clinical-grade theragnostic biomarkers.

## MATERIALS AND METHODS

### Study design

The objective of this study was to determine whether cryptic exons generate *de novo* peptides in the setting of TDP-43 deficiency. To develop a deep and high-quality neuronal cryptic exon catalog, we performed total RNA-seq and differential splicing analyses on CRISPRi TDP-43 depleted human iPSC neurons. We then designed an unbiased proteogenomic pipeline, using parallel RNA-seq and shotgun proteomics in TDP-43 KD iPSC neurons, to identify peptides that map to CE coordinates. Next, we cross-referenced our iPSC neuron-predicted CEs against datasets from FTD/ALS post-mortem brain tissue to test the fidelity of our *in vitro* model in predicting TDP-43 pathology in human brains. We used western blot and targeted proteomics to validate the presence of *de novo* peptides in TDP-43 KD iPSC neurons. We then examined the impact of *de novo* peptides on the biology of the proteins in which they are expressed via affinity purification mass spectrometry. Finally, we developed a novel targeted proteomics assay to demonstrate the presence of *de novo* “cryptic” peptides in the CSF of ALS patients. Sample size was determined empirically in accordance with field standards to ensure sufficient power for detecting statistical differences. The number of experimental replicates is provided in each figure legend. Only RNA and protein of low abundance or quality were excluded. All ALS CSF samples that were provided to us from the Mayo Clinic were analyzed, and none were excluded.

### CRISPRi knockdown experiments

CRISPRi knockdown experiments were performed in the WTC11 iPSC line harboring stable TO-NGN2 and dCas9-BFP-KRAB cassettes at safe harbor loci (*35*). CRISPRi knockdown of TDP-43 was achieved by infecting iPSCs with a lentiviral-expressed sgRNA targeting the transcription start site of TARDBP (or non-targeting control sgRNA) (*13*). Neurons were differentiated as described previously (*36, 37*) and harvested 17 days post-differentiation for all experiments. See supplemental methods section for additional details.

### Short-read RNA sequencing, differential gene expression, and splicing analysis

RNA was extracted from day 17 neurons using the Direct-zol RNA Miniprep Kit (Zymo Research R2051) and sequenced on a Novaseq 6000 (2 x 150bp paired-end). Sequencing files were trimmed for adapters (cutadapt *v2.5*)(*38*), quality checked (FastQC *v0.11.6*)(*39*), and aligned to GRCh38 reference genome (STAR *v2.7.3a*)(*40*). Differential expression analysis was performed using the standard DESeq2 workflow(*41*). For differential splicing analysis, all samples were run through MAJIQ (*42*) in the same manner using a custom Snakemake pipeline (https://github.com/frattalab/splicing). An additional pipeline was developed to visualize and categorize each mis-spliced junction as cryptic exon, exon skip, intron retention, or canonical junction (https://github.com/NIH-CARD/proteogenomic-pipeline). See supplemental methods section for additional details.

### Long-read sequencing and data processing

Sequence-specific and regular cDNA-PCR libraries were prepared using the Oxford Nanopore Technologies sequencing kit (SQK-PCS109) and sequenced on a Promethion device. Sequencing data were basecalled (Guppy v3.4.5) and mapped to GRCh38 reference genome (minimap2)(*43*). Basecalled reads were also independently aligned against the human transcriptome (Ensembl version 92). Transcript identification and quantification was performed with Bambu at maximum sensitivity. Only reads overlapping mis-spliced genes were used. See supplemental methods section for additional details.

### Ribosome profiling

Ribosome profiling libraries, from three biological replicates both of control and TDP-43 knockdown in iPSC neurons, were derived from a previous study, but underwent Cas9-based rRNA depletion followed by resequencing on an Illumina Hi-Seq 4000 machine to improve read depth (*13, 44*). Multi-mapping reads were discarded and reads 28-30 nt in length were selected for analysis. See supplemental methods section for additional details.

### *De novo* peptide sequence prediction

Dasper was used to classify each mis-spliced junction as novel acceptor, novel donor, or exon skipping event(*45*). Exonic regions were identified using Gencode v31 reference annotation and two different transcript assembly tools, Scallop (*46*) and Stringtie2(*47*). All novel exonic regions were mapped back to coordinates that overlapped cryptic junctions. Transcripts overlapping with cryptic events were identified as “backbone transcripts” and used to insert the novel cryptic event. The resulting genomic regions were used to extract nucleotide sequences. For each nucleotide sequence, the amino acid sequence of all possible open reading frames (ORF) between every methionine and stop codon was extracted. See supplemental methods section for additional details.

### Liquid chromatography and mass spectrometry

Protein tryptic digestion was performed using a fully automated sample preparation workflow as previously described (*48*). For human CSF samples, the top 14 most abundant proteins were depleted with High Select Depletion spin column and resin (Thermo Scientific A36369). Three data acquisition approaches (DDA, DIA, and PRM) were used in LC-MS/MS analyses. iNeuron samples were separated on a Waters AQUITY UPLC M_Class System and injected into an Orbitrap Exploris 480 for DIA proteomics. All other samples were separated on an Ultimate 3000 nano-LC system and injected into a high-resolution Orbitrap Eclipse MS. Protein tryptic digestion was performed using a fully automated sample preparation workflow as previously described (*48*). For human CSF samples, the top 14 most abundant proteins were depleted with High Select Depletion spin column and resin (Thermo Scientific A36369). Three data acquisition approaches (DDA, DIA, and PRM) were used in LC-MS/MS analyses. See supplemental methods section for additional details.

### Proteomics database search and statistical analysis

DDA-based discovery proteomics was performed using a custom database of *de novo* peptides (File S1) in PEAKS studio v10.6 (*49*). Skyline software (*50*) was used for quantification and visualization of PRM and DIA targeted proteomics data. Statistical analyses were conducted in R studio, using a two-sided t-test and Benjamini-Hochberg adjustment for multiple comparisons. Differential expression analysis was carried out in the Spectronaut software (*51*), generating log fold changes and q-values for volcano plot analysis. See supplemental methods section for additional details.

### Splicing analysis of postmortem brain tissue

Data from FACS-sorted frontal cortex neuronal nuclei were obtained from the Gene Expression Ombius (GEO) database (<u>GSE126543</u>) and aligned to the GRCh38 reference genome as previously described (*13*). The splice junction output tables were then clustered and converted into PSI metrics using Dasper (*45*). As splicing tools can be prone to one-off errors for exact splice junction coordinates, the 340 *bona fide* splicing events from MAJIQ were manually curated against the automated splice junctions from STAR to confirm the absence or presence of each event in the FACs-sorted nuclei.

The analysis of postmortem brain tissue from NYGC contains 472 neurological tissue samples from 286 individuals in the NYGC ALS dataset, including non-neurological disease controls, FTLD, ALS, FTD with ALS (ALS-FTLD), and ALS with suspected Alzheimer’s disease (ALS-AD). The NYGC dataset was analyzed for the 340 *bona fide* splicing events from MAJIQ using a modified splice junction parsing pipeline. Junctions from NYGC samples that overlapped with the 340 events were manually curated to ensure accuracy. Read counts for each splice junction were normalized and converted to z-scores to calculate AUC scores for TDP-43 pathology classification performance using the pROC package (*52*). Meta-scores were created using z-scores across all cryptic junctions, junctions with a positive predictive value above 0.6, and predictive junctions excluding STMN2. See supplemental methods section for additional details.

### qRT-PCR validation of cryptic exons

Quantitative real-time PCR (qRT-PCR) was conducted using SYBR GreenER qPCR SuperMix (Invitrogen) for all samples in triplicates. qRT-PCR were run in a QuantStudio™ 7 Flex Real-Time PCR System (Applied Biosystems). Relative quantification was determined using the ΔΔCt method and normalized to the endogenous controls *GAPDH* and *RPLP0*. Primer efficiency was verified for each cryptic exon prior to running the qRT-PCRs. See Table S3E for list of primers. Additional details are provided in the supplemental methods section.

### Western blot

MYO18A-CE antibody was generated by Labcorp by immunizing rabbits with a peptide including the complete 20 residue neoepitope (VKEEDKTLPKPGSPGKEEGA). Equal amounts of protein were loaded on 3-8% Tris-Acetate (MYO18A-CE) or 4-20% Tris-Glycine (TDP-43 and GAPDH) gels and transferred to membranes. The primary antibodies used for the MYO18A-CE experiment were anti-rabbit MYO18A-CE antibody (1:500), anti-rabbit TDP-43 antibody (1:1000, Proteintech, 12892-1-AP), and anti-mouse GAPDH antibody (1:5000, Meridian Life Science, H86504M). For analysis of total HDGFL2 protein levels, equal amounts of protein were loaded in 7% Bis-Tris gels and run with MOPS buffer (Thermo Scientific) and transferred to membranes (Amersham GE Healthcare). Primary antibodies used were rabbit anti-HDGFL2 (1:1000 Sigma HPA 044208), mouse anti-TDP-43 (1:5000 Abcam ab104223) and rat ant-Tubulin (1:5000 Millipore MAB1864). Western blots were developed on a Bio-Rad ChemiDoc and quantified with ImageJ. See supplemental methods section for additional details.

### Affinity purification mass spectrometry

Anti-IgG or anti-HDGFL2-CE antibodies were added to cell lysates and incubated overnight. Sample protein concentrations were evaluated using a detergent compatible protein assay (DCA) (Bio-Rad, Hercules, CA, Cat. #5000111). Automated affinity purification mass spec was performed using an automated protocol on the KingFisher Flex Purification System, followed by overnight incubation. Peptides were dried and reconstituted in a 2% acetonitrile and 0.4% trifluoroacetic acid (TFA) solution. Peptide concentrations were evaluated on a Denovix DS-11 FX Spectrophotometer/Fluorometer and normalized for analysis by LC-MS/MS. Precursor matching, protein inference, and quantification were performed in the Spectronaut software (*51*) using the DirectDIA workflow in default settings. Differential abundance analysis was carried out in the Spectronaut software 16.2 (*51*), generating log fold changes and q-values by using a two-sided t-test and Benjamini-Hochberg adjustment for multiple comparisons. R software version 4.2.3 was used for data visualization, and ShinyGo was used for gene ontology analysis. See supplemental methods section for additional details.

### STRING analysis

Gene ontology analysis was performed on significantly enriched HDGFL2 interacting proteins, compared to control IgG pulldowns, or HDGFL2 interacting proteins that were significantly changed in their interactions depending on TDP-43 knockdown. GO term analysis was performed in ShinyGo v0.76.3(*53*). String analysis was performed using String v11.5 (*54*) and resulting networks were downloaded and visualized using Cytoscape (v3.9.1) (*55*).

### CSF samples

Study samples were collected through the Neurological Disease Biorepository and Biomarker Initiative at Mayo Clinic’s campus in Jacksonville, Florida (IRB#13-004314 and IRB# 11-002986). Our cohort of cases consisted of 9 males and 6 females (all white race and not of Hispanic ethnicity), with median age at CSF collection of 64 years (range: 50-78) and median age of ALS onset of 59 years (range: 49-79 years).

### Statistical analysis

Raw, replicate-level data for each experiment is provided in the supplementary materials. Graphs were generated using GraphPad Prism Version 9.3.1. Data are displayed as mean ± SEM; individual data points are also shown. Statistical testing was performed using GraphPad Prism Version 9.3.1 and R statistical programming language. Shapiro-Wilk test was used to assess normality. *P*-values for normally distributed datasets were determined by unpaired *t*-test, with Benjamini-Holchberg adjustment for multiple comparisons. For non-normally distributed data, nonparametric analysis was carried out using the Mann-Whitney test or one-way ANOVA. *p* < 0.05 was considered significant for two-tailed tests, while *p* <0.1 was considered significant for one-tailed tests. The number of biological replicates and details of statistical analyses are provided in the figure legends.

## List of Supplementary Materials

**Supplementary Fig. S1.** TDP-43 downregulation in iPSC neurons causes mis-splicing and loss of associated transcript and protein products.

**Supplementary Fig. S2.** Identification of full-length transcripts expressing cryptic peptides with Nanopore long-read sequencing

**Supplementary Fig. S3.** Validation of cryptic RNA enrichment in TDP-43 deficient iPSC neurons

**Supplementary Fig. S4.** Cryptic exon inclusion alters the interacting partners of affected proteins

**Supplementary Fig. S5.** Cryptic peptide validation in TDP-43 deficient iPSC neurons via targeted proteomics

**Supplementary Fig. S6.** Design of targeted proteomics assay for cryptic peptide detection in ALS CSF

**Supplementary Fig. S7.** Detection of cryptic peptides in ALS CSF

## Supplementary Methods

**Supplementary File S1.** Custom database of *de novo* peptides

**Supplementary File S2.** Sashimi plots of mis-spliced junctions in TDP-43 KD iNeurons

**Supplementary Table S1A.** Differentially spliced junctions in TDP-43 KD vs control iPSC neurons

**Supplementary Table S1B.** Differential ribosomal profiling in TDP-43 KD vs control iPSC neurons

**Supplementary Table S1C.** Differential transcript abundance in TDP-43 KD vs control iPSC neurons

**Supplementary Table S1D.** Differential protein abundance in TDP-43 KD vs control iPSC neurons

**Supplementary Table S2A.** Shotgun proteomics database search results

**Supplementary Table S2B.** Amino acid sequence of putative cryptic peptides across 12 genes

**Supplementary Table S2C.** List of targeted long-read sequencing primers

**Supplementary Table S3A.** Lookup table of mis-spliced junctions and corresponding symbols

**Supplementary Table S3B.** Percent spliced in (PSI) values of iPSC neuron predicted cryptic exons in FACS sorted TDP-43 positive and negative neuronal nuclei from postmortem FTD/ALS cortex

**Supplementary Table S3C.** AUC of iPSC neuron-predicted cryptic exons in postmortem bulk RNAseq from FTLD temporal cortex

**Supplementary Table S3D.** AUC of iPSC neuron-predicted cryptic exons in postmortem bulk RNAseq from ALS motor cortex

**Supplementary Table S3E.** QRTPCR primer sequences for cryptic exons validated in FTLD-TDP frontal cortex

**Supplementary Table S4A.** List of co-immunoprecipitated proteins using anti-HDGFL2 antibody vs control IgG.

**Supplementary Table S4B.** Affinity purification mass spectrometry analysis of protein-protein interactions in TDP-43 KD vs control iPSC neurons

**Supplementary Table S5A.** Lookup table of cryptic peptide amino acid sequences and corresponding symbols

**Supplementary Table S5B.** PRM intensity values of cryptic peptides in TDP-43 KD and control iPSC neurons

**Supplementary Table S6A.** Trait file for the C9+ vs C9-ALS samples in the TMT-MS assay

**Supplementary Table S6B.** Ranked protein intensities in CSF from C9+ vs C9-ALS samples

**Supplementary Table S6C.** Ranked protein intensities in CSF from pooled ALS samples

**Supplementary Table S6D.** Ranked protein intensities in CSF from a published dataset (*56*)

**Supplementary Table S6E.** ALS CSF proteins vs iPSC neuron-predicted cryptic exons

**Supplementary Table S6F.** Heatmap intensity values for cryptic peptides detected in ALS CSF

**Supplementary Table S6G.** DIA-targeted proteomics intensity values for all cryptic peptides detected in ALS CSF

## Acknowledgements

We are grateful for the thoughtful input provided by Andy Singleton, Mark Cookson, and Caroline Pantazis. We thank Mandy Guo for her assistance with CSF experiments and gratefully acknowledge support from the FLI Core Facility Proteomics.

## Author contributions

Conceptualization: SS, YAQ, ALB, LP, PF, MEW
Methodology: SS, YAQ, ALB, CB, OGW, CC, AO, LP, PF, MEW Investigation: SS, YAQ, ALB, OGW, CB, CB, YZ, MP, MJK, AK, SP, SEH, JH, DMR, HY, JR, EKS, SIS, JCM, JFR, VHR, MPN, ZL, AS, LP, DMD, MS, NS, BO, YK
Visualization: SS, YAQ, ALB, OGW, SIS, MAN, MP, MEW
Funding acquisition: JU, MP, LP, PF, MEW
Project administration: LS, AA, JR, AM, LP, PF, MEW
Supervision: YAQ, MP, HY, MAN, JYK, SJ, JDG, AO, NTS, MM, LP, PF, MEW
Writing – original draft: SS, ALB, PF, MEW
Writing – review & editing: SS, YAQ, ALB, OGW, CB, CB, YZ, MP, MJK, DMR, JR, EKS, SIS, MAN, LP, DMD, AO, NTS, MM, LP, PF, MEW

## Data and materials availability

Analysis code and data to reproduce all figures are available at: https://github.com/NIH-CARD/proteogenomic-pipeline. RNA-seq and proteomics data files can be accessed at Alzheimer’s Disease Workbench (ADWB): https://www.alzheimersdata.org/ad-workbench. ALS C9+/- CSF TMT data can be downloaded from Synapse (SynID: syn25795030): https://www.synapse.org. Long-read sequencing data are available on The NHGRI Analysis Visualization and Informatics Lab-space (AnVIL): https://anvilproject.org/

