## Supplementary Materials for "Mis-spliced transcripts generate *de novo* proteins in TDP-43-related ALS/FTD"

**List of Supplementary Materials**

*Please note: supplementary figures and methods are provided below; supplementary files and tables can be accessed here (https://shorturl.at/HJORU)*

**Supplementary Fig. S1.** TDP-43 downregulation in iPSC neurons causes mis-splicing and loss of associated transcript and protein products.

**Supplementary Fig. S2.** Identification of full-length transcripts expressing cryptic peptides with Nanopore long-read sequencing

**Supplementary Fig. S3.** Validation of cryptic RNA enrichment in TDP-43 deficient iPSC neurons

**Supplementary Fig. S4.** Cryptic exon inclusion alters the interacting partners of affected proteins

**Supplementary Fig. S5.** Cryptic peptide validation in TDP-43 deficient iPSC neurons via targeted proteomics

**Supplementary Fig. S6.** Design of targeted proteomics assay for cryptic peptide detection in ALS CSF

**Supplementary Fig. S7.** Detection of cryptic peptides in ALS CSF

**Supplementary Methods**

**Supplementary File S1.** Custom database of *de novo* peptides

**Supplementary File S2.** Sashimi plots of mis-spliced junctions in TDP-43 KD iNeurons

**Supplementary Table S1A.** Differentially spliced junctions in TDP-43 KD vs control iPSC neurons

**Supplementary Table S1B.** Differential ribosomal profiling in TDP-43 KD vs control iPSC neurons

**Supplementary Table S1C.** Differential transcript abundance in TDP-43 KD vs control iPSC neurons

**Supplementary Table S1D.** Differential protein abundance in TDP-43 KD vs control iPSC neurons

**Supplementary Table S2A.** Shotgun proteomics database search results

**Supplementary Table S2B.** Amino acid sequence of putative cryptic peptides across 12 genes

**Supplementary Table S2C.** List of targeted long-read sequencing primers

**Supplementary Table S3A.** Lookup table of mis-spliced junctions and corresponding symbols

**Supplementary Table S3B.** Percent spliced in (PSI) values of iPSC neuron predicted cryptic exons in FACS sorted TDP-43 positive and negative neuronal nuclei from postmortem FTD/ALS cortex

**Supplementary Table S3C.** AUC of iPSC neuron-predicted cryptic exons in postmortem bulk RNAseq from FTLD temporal cortex

**Supplementary Table S3D.** AUC of iPSC neuron-predicted cryptic exons in postmortem bulk RNAseq from ALS motor cortex

**Supplementary Table S3E.** QRTPCR primer sequences for cryptic exons validated in FTLD-TDP frontal cortex

**Supplementary Table S4A.** List of co-immunoprecipitated proteins using anti-HDGFL2 antibody vs control IgG.

**Supplementary Table S4B.** Affinity purification mass spectrometry analysis of protein-protein interactions in TDP-43 KD vs control iPSC neurons

**Supplementary Table S5A.** Lookup table of cryptic peptide amino acid sequences and corresponding symbols

**Supplementary Table S5B.** PRM intensity values of cryptic peptides in TDP-43 KD and control iPSC neurons

**Supplementary Table S6A.** Trait file for the C9+ vs C9- ALS samples in the TMT-MS assay

**Supplementary Table S6B.** Ranked protein intensities in CSF from C9+ vs C9- ALS samples

**Supplementary Table S6C.** Ranked protein intensities in CSF from pooled ALS samples

**Supplementary Table S6D.** Ranked protein intensities in CSF from a published dataset *(56)*

**Supplementary Table S6E.** ALS CSF proteins vs iPSC neuron-predicted cryptic exons

**Supplementary Table S6F.** Heatmap intensity values for cryptic peptides detected in ALS CSF

**Supplementary Table S6G.** DIA-targeted proteomics intensity values for all cryptic peptides detected in ALS CSF

### Fig S1

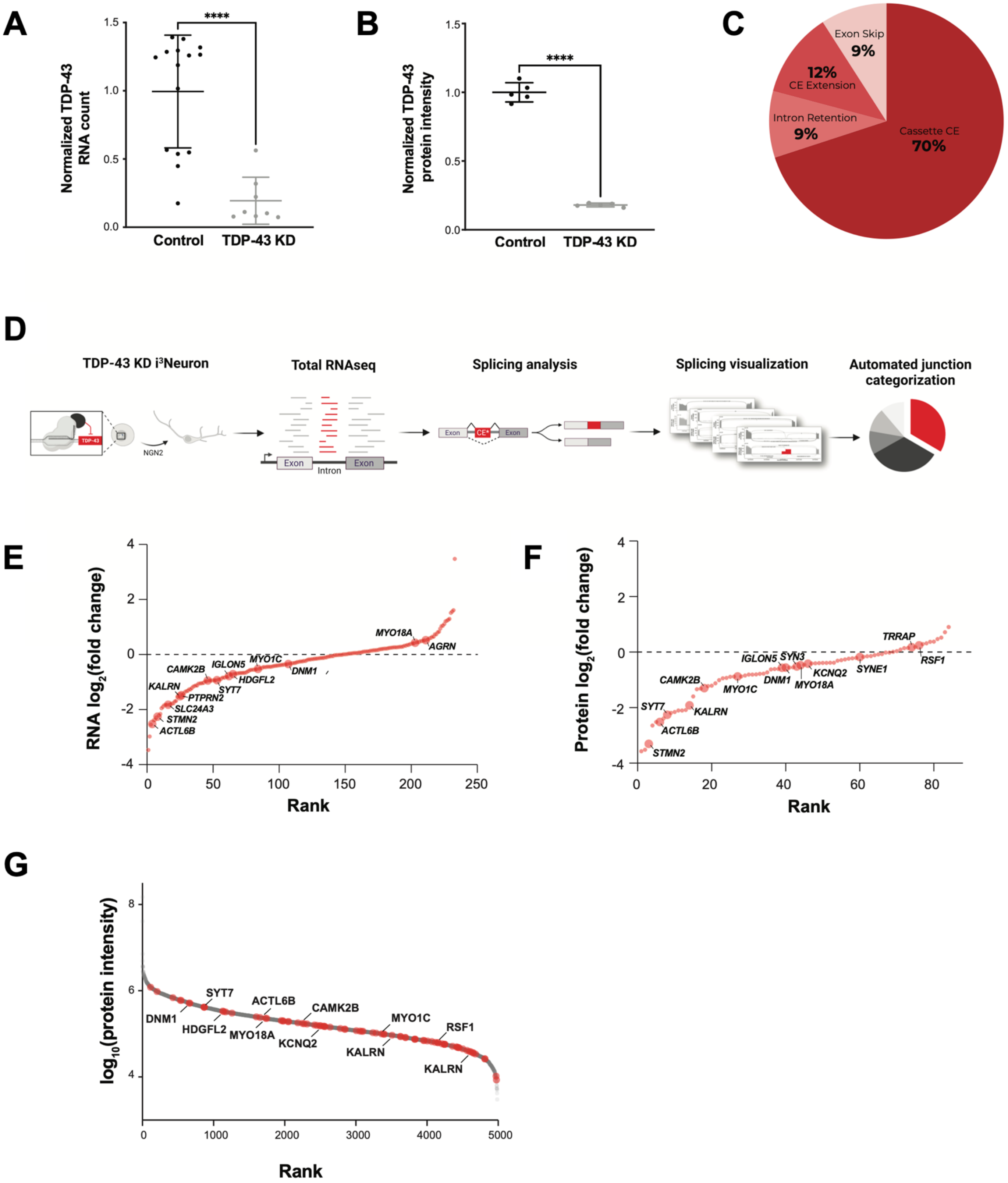

A
Normalized TDP-43 RNA count
Control
TDP-43 KD
B
Normalized TDP-43 protein intensity
C
Exon Skip 9%
12% CE Extension
Intron Retention 9%
Cassette CE 70%
D
TDP-43 KD i³Neuron
Total RNAseq
Splicing analysis
Splicing visualization
Automated junction categorization
E
RNA log₂(fold change)
Rank
F
Protein log₂(fold change)
Rank
G
log₁₀(protein intensity)
Rank

**Supplementary Fig. S1. TDP-43 downregulation in iNeurons causes mis-splicing and loss of associated transcript and protein products.** (**A**) Differential TDP-43 transcript abundance in control and TDP-43 KD iPSC neurons (iNeurons), quantified using total short-read RNAseq. P-value shown from one-way t-test. **** $p<0.0001$(**B**) Differential TDP-43 protein abundance in control and TDP-43 KD iNeurons, quantified using DIA total proteomics. P-value shown from one-way t-test. **** $p<0.0001$ (**C**) Mis-spliced junction types in TDP-43 KD iNeurons (**D**) Proteogenomic workflow to identify and characterize TDP-43 related mis-spliced junctions in iNeurons (**E**) Rank plot of LFC in cryptic exon RNA abundance in control vs TDP-43 KD iNeurons (**F**) Rank plot of LFC in cryptic exon parent protein abundance in control vs TDP-43 KD iNeurons (**G**) Rank plot of baseline protein abundance in iNeurons. Parent proteins of cryptic exon-harboring genes are shown in red.

### Fig S2

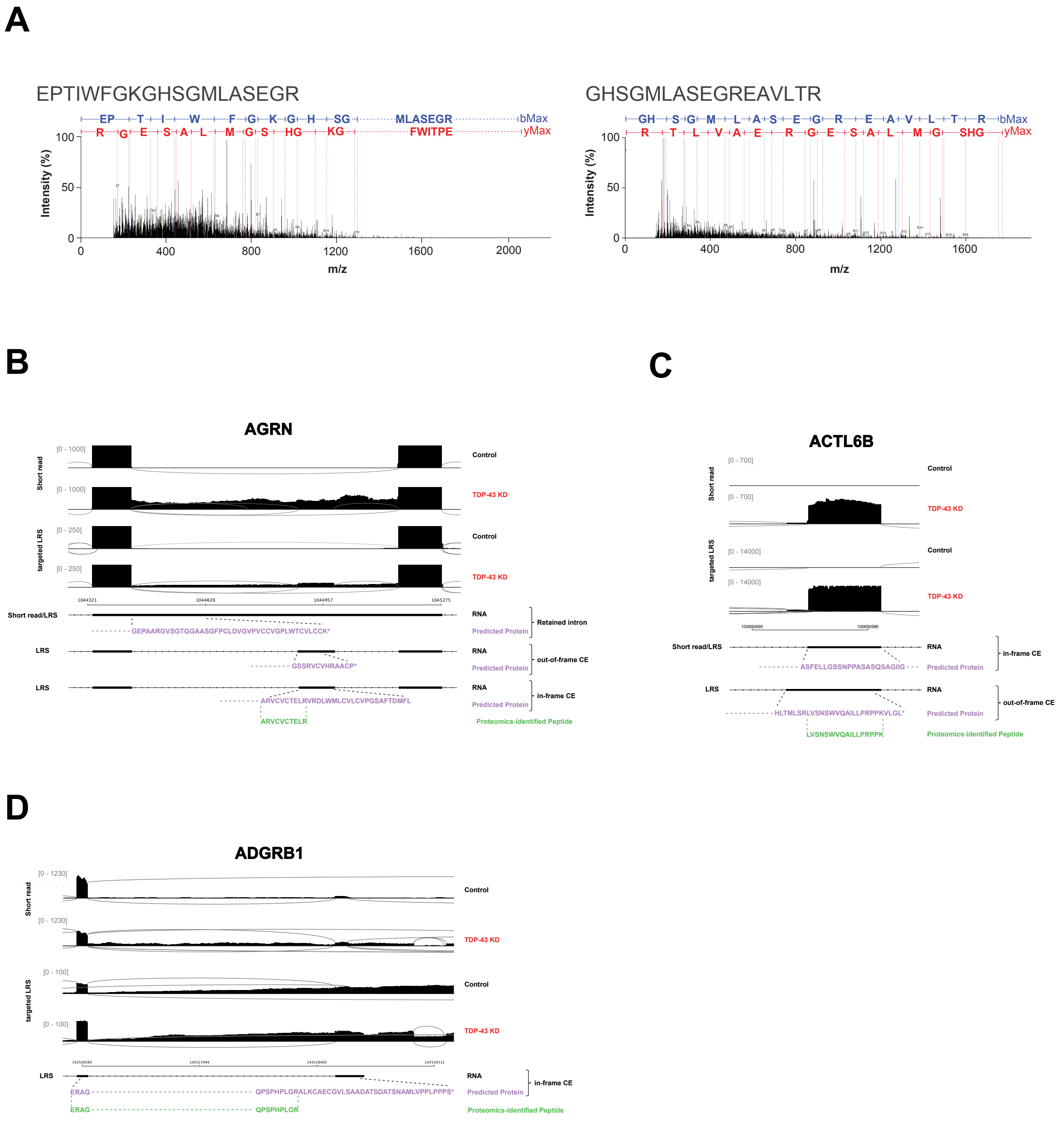
A
EPTIWFGKGHSGMLASEGR
GHSGMLASEGREAVLTR
bMax
yMax
Intensity (%)
m/z
B
AGRN
Short read
targeted LRS
Control
TDP-43 KD
Short read/LRS
LRS
RNA
Predicted Protein
Retained intron
out-of-frame CE
in-frame CE
GEPAARGVSGTGGAASGFPCLDVGVPVCCVGPLWTCVLCCK*
GSSRVCVHRAACP*
ARVCVCTELRVRDLWMLCVLCVPGSAFTDMFL
ARVCVCTELR
Proteomics-identified Peptide
C
ACTL6B
ASFELLGSSNPPASASQSAGIIG
HLTMLSRLVSNSWVQAILLPRPPKVLGL*
LVSNSWVQAILLPRPPK
D
ADGRB1
ERAG
QPSPHPLGRALKCAECGVLSAADATSDATSNAMLVPPLPPPS*
QPSPHPLGR

**Supplementary Fig. S2. Identification of full-length transcripts expressing cryptic peptides with Nanopore long-read sequencing.** (**A**) MS/MS spectra of two cryptic peptides that map to the HDGFL2 cryptic exon, identified using label-free DIA mass spectrometry. (**B-D**) Representative sashimi plots comparing Illumina short-read RNAseq and Nanopore long-read RNAseq for *AGRN* (B), *ACTL6B* (C) and *ADGRB1* (D) transcripts in control and TDP43-KD iNeurons. The corresponding transcripts (black) identified by Nanopore long-read sequencing, their predicted amino-acid sequences (purple), and cryptic peptides (green) are shown. Transcripts were identified with bambu.

Fig S3

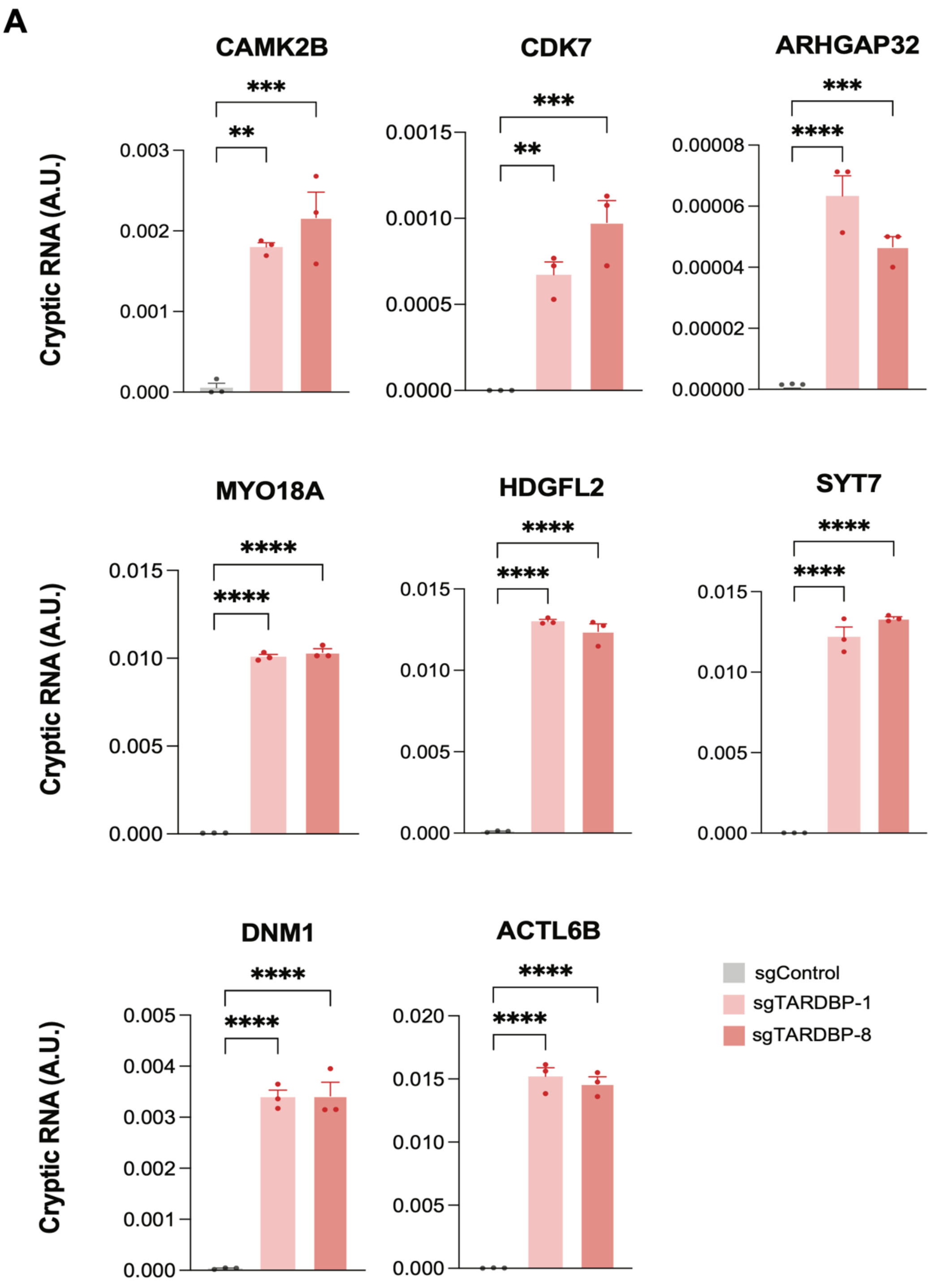
A
CAMK2B
CDK7
ARHGAP32
MYO18A
HDGFL2
SYT7
DNM1
ACTL6B
Cryptic RNA (A.U.)
sgControl
sgTARDBP-1
sgTARDBP-8

**Supplementary Fig. S3. Validation of cryptic RNA enrichment in TDP-43 deficient iNeurons.** (**A**) qRT-PCR analyses of RNA from iNeurons treated with control sgRNA (sgControl) or two different guides against *TARDBP* (sgTARDBP-1, sgTARDBP-8), which lead to ~50% reduction in *TARDBP* RNA and TDP-43 protein levels. Data are presented as mean +/- SEM. *P*-values shown are from one-way ANOVA: *$p \leq 0.05$, **$p \leq 0.005$, *** $p \leq 0.0005$; ****$p \leq 0.0001$.

### Fig S4

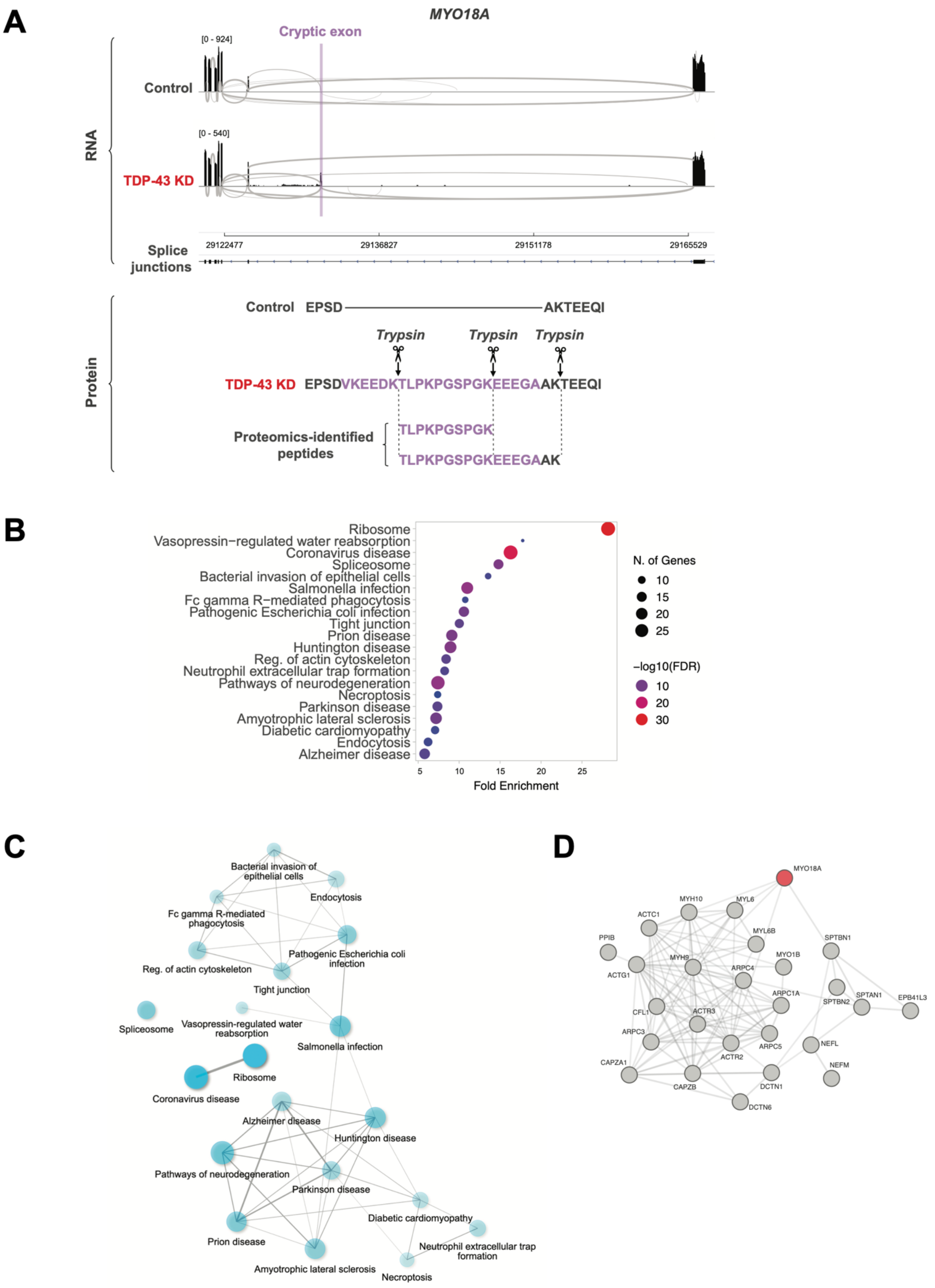
A
MYO18A
Cryptic exon
[0 - 924]
Control
[0 - 540]
TDP-43 KD
RNA
29122477
29136827
29151178
29165529
Splice junctions
Protein
Control EPSD AKTEEQI
Trypsin
Trypsin
Trypsin
TDP-43 KD EPSDVKEEDKTLPKPGSPGKEEEGAAKTEEQI
Proteomics-identified peptides
TLPKPGSPGK
TLPKPGSPGKEEEGAAK
B
Ribosome
Vasopressin−regulated water reabsorption
Coronavirus disease
Spliceosome
Bacterial invasion of epithelial cells
Salmonella infection
Fc gamma R−mediated phagocytosis
Pathogenic Escherichia coli infection
Tight junction
Prion disease
Huntington disease
Reg. of actin cytoskeleton
Neutrophil extracellular trap formation
Pathways of neurodegeneration
Necroptosis
Parkinson disease
Amyotrophic lateral sclerosis
Diabetic cardiomyopathy
Endocytosis
Alzheimer disease
5
10
15
20
25
Fold Enrichment
N. of Genes
10
15
20
25
−log10(FDR)
10
20
30
C
Bacterial invasion of epithelial cells
Endocytosis
Fc gamma R-mediated phagocytosis
Pathogenic Escherichia coli infection
Reg. of actin cytoskeleton
Tight junction
Spliceosome
Vasopressin-regulated water reabsorption
Salmonella infection
Ribosome
Coronavirus disease
Alzheimer disease
Huntington disease
Pathways of neurodegeneration
Parkinson disease
Diabetic cardiomyopathy
Neutrophil extracellular trap formation
Prion disease
Amyotrophic lateral sclerosis
Necroptosis
D
MYO18A
MYH10
MYL6
ACTC1
PPIB
MYL6B
SPTBN1
MYH9
MYO1B
ACTG1
ARPC4
ARPC1A
SPTAN1
EPB41L3
SPTBN2
CFL1
ACTR3
ARPC3
ARPC5
NEFL
ACTR2
CAPZA1
CAPZB
DCTN1
NEFM
DCTN6

**Supplementary Fig. S4. Cryptic exon inclusion alters the interacting partners of affected proteins.** (**A**) TDP-43 loss causes formation of a cryptic exon in MYO18A (sashimi plot shown, top), with a predicted in-frame cryptic exon (purple). Two cryptic peptides, detected by proteogenomics, map to this in-frame cryptic exon (bottom). (**B**) KEGG pathway analysis of HDGFL2 interacting proteins in iNeurons that were identified via DIA APMS (anti-HDGFL2 vs control IgG pulldown). (**C**) Graph representation of HDGFL2-interacting proteins associated with the KEGG pathways identified in (B). (**D**) STRING diagram of the subset of HDGFL2 interacting proteins associated with MYO18A interactions and cytoskeleton regulation.

### Fig S5

## A

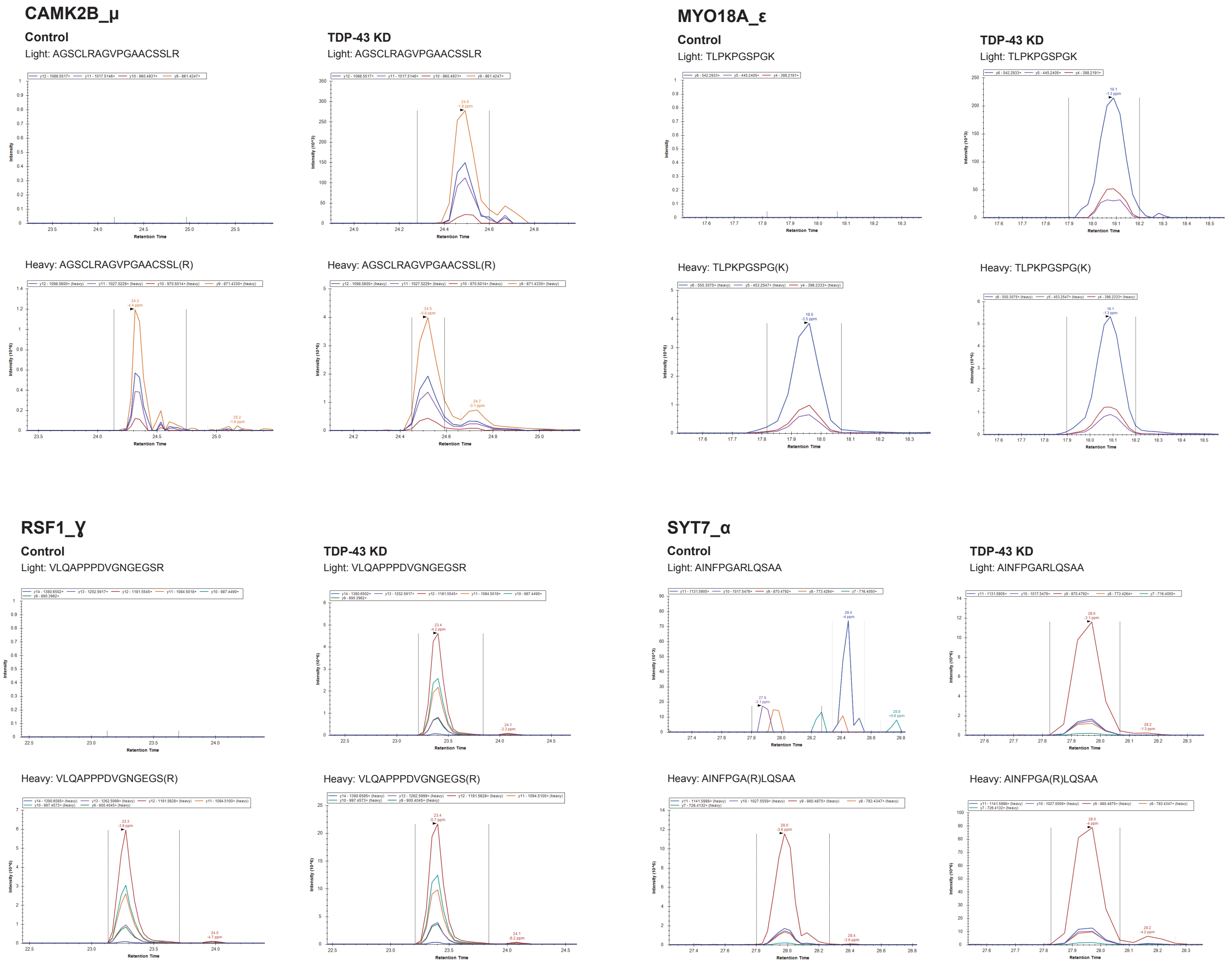


**Supplementary Fig. S5. Cryptic peptide validation in TDP-43 deficient iPSC neurons via targeted proteomics.** (A) Spectral plot of heavy standards and light (endogenous) ions for 4 representative cryptic peptides in TDP-43 KD and control iNeurons.

#### Fig S6

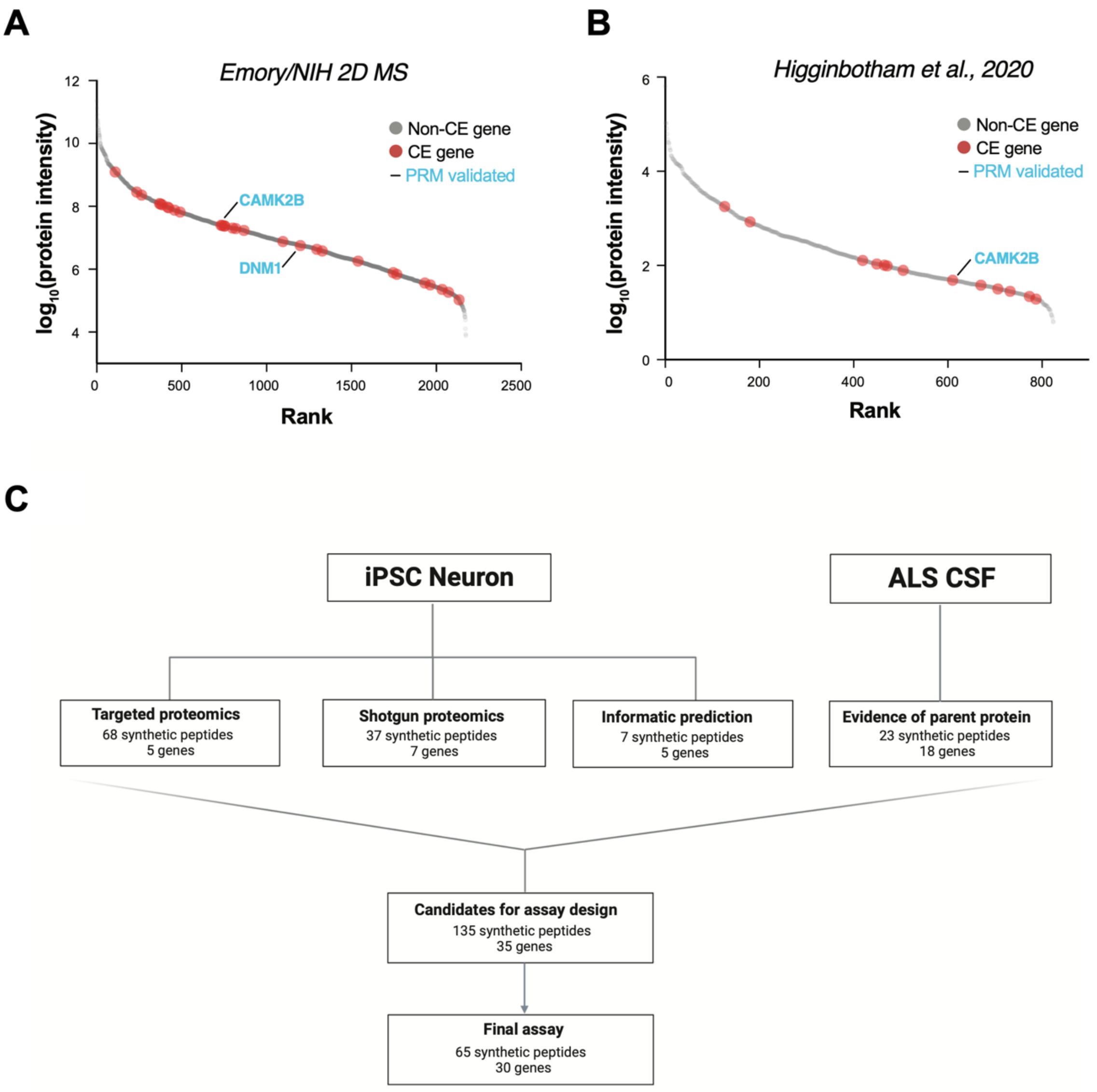


**Supplementary Fig. S6. Design of targeted proteomics assay for cryptic peptide detection in ALS CSF.** (**A-B**) Rank plot of proteins detected in CSF from patients with ALS from an internal (A) and previously published (B) dataset. Parent proteins of iPSC neuron-predicted cryptic exon genes are shown in red. (**C**) Selection process for heavy isotope-labeled peptides for use in targeted proteomics assay

### Fig S7

## A

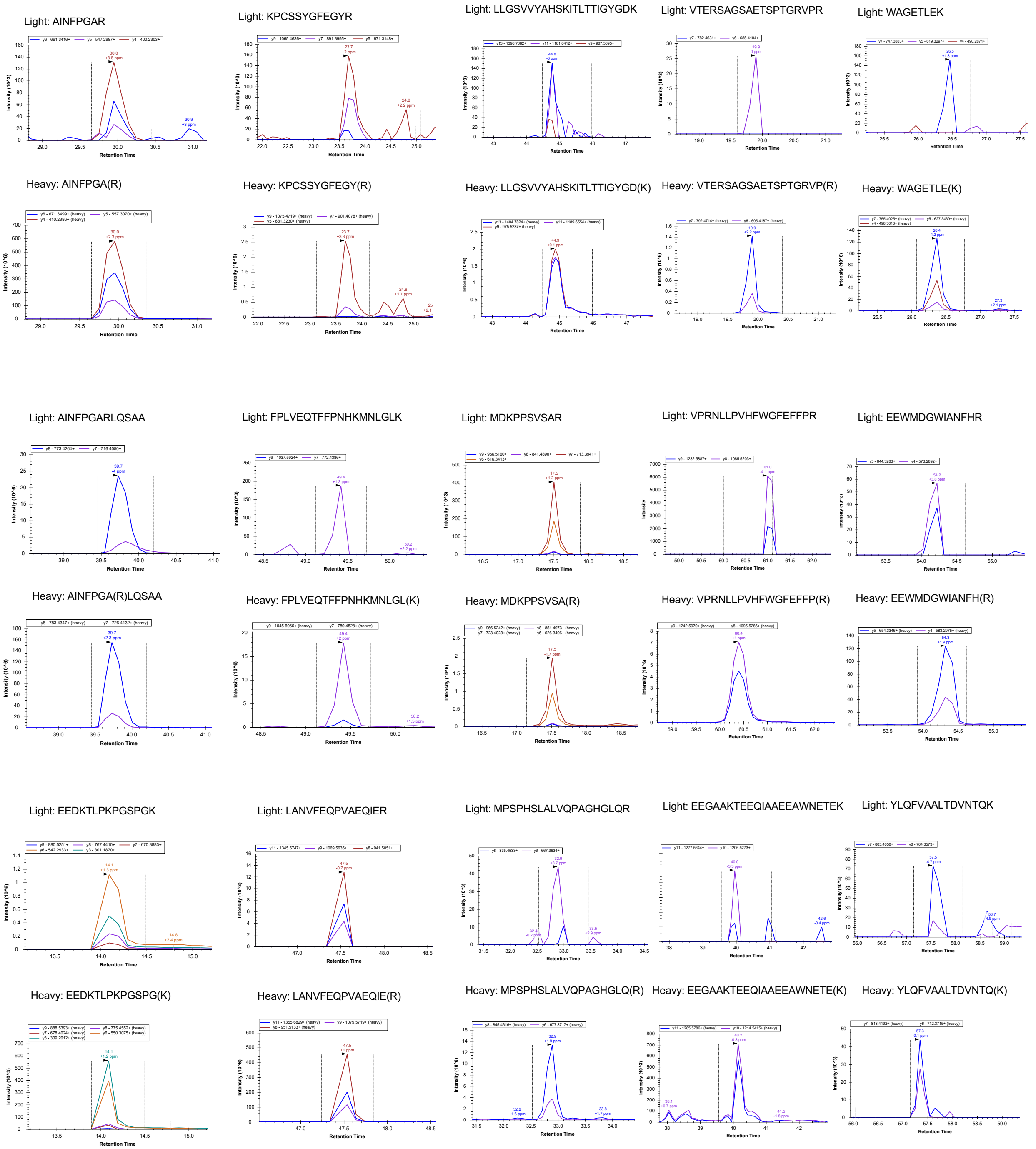


**Supplementary Fig. S7. Detection of cryptic peptides in ALS CSF.** (**A**) Spectral plot of heavy standards and light (endogenous) ions for cryptic peptides in representative ALS CSF samples.

#### Supplementary Methods

#### CRISPRi knockdown experiments

##### *Cell culture maintenance*

Human iPSCs were grown as previously described *(36, 37)*. Briefly, cells were grown on Matrigel (Corning # 354277)-coated cell culture plates in Essential 8 media (ThermoFisher Scientific # A1517001). Cells were dissociated by incubation with Accutase (Life Technologies # A1110501) for 5 minutes at 37°C and pelleted by centrifugation at 200 x g for 5 minutes. Cell pellets were washed with PBS and pelleted again by centrifugation at 200 x g for 5 minutes. Final pellets were resuspended in E8 supplemented with the ROCK inhibitor Y-27632 (Selleckchem # S1049) to promote adherence and survival and plated onto Matrigel-coated cell culture plates. Media was changed every 1-2 days and Y-27632 was removed from cell culture media when cells formed colonies of > 20 cells.

##### *CRISPRi-mediated gene knockdown in human iPSCs*

CRISPRi knockdown experiments were performed in the i11w-mNC line, a derivative of the previously described WTC11 line *(13, 35)*. This line stable expresses CAG-dCas9-BFP-KRAB at the CLYBL safe harbor locus and a tetracycline-inducible mNGN2 transgene at the AAVS1 locus. sgRNAs targeting TARDBP or a non-targeting control sgRNA were delivered by iPSCs via lentiviral transduction. iPSCs were split and treated with lentivirus by resuspension of the final cell pellet in PBS containing lentivirus (day -4). Cells were incubated in PBS-lentivirus for 5 min and added dropwise to cell culture plates containing E8 + 10 μM Y-27632 media. 24 hours post-transduction (day -3), cells were washed with PBS and fresh E8 media was added (without Y-27632). On day -2, cells were dissociated and replated in E8 media containing 10 μM Y-27632 and 10ug/ml puromycin to select for cells that were successfully transduced with the sgRNA lentivirus. On day -1, cells were washed with PBS to remove debris, and fresh E8 + puromycin (10µg/ml final concentration) was added to the cells. On day 0, cells were visually inspected and >95% BFP+, suggesting >95% of cells expressed sgRNA. Cells were replated using the above method and plated into Neuronal Induction Media containing Knockout DMEM/F12 media (Life Technologies # 12660012), N2 supplement (Life Technologies # 17502048), 1× GlutaMAX (ThermoFisher Scientific # 35050061), 1× MEM Non-essential Amino Acids (Thermofisher Scientific # 11140050), 10 µM Y-27632 and 2 µg/ml doxycycline (Clontech # 631311). On day 1 and day 2, media was changed, and fresh Neuronal Induction Media was added, excluding Y-27632. On day 3, cells were dissociated and replated using the above method, and replated into poly-L-ornithine (0.1 mg/ml; Sigma # P3655-10MG)-coated cell culture plates at a density of 1.5 million cells per well of a 6-well plate. From day 3 until the end of the experiment (day 17), cells were cultured in Neuronal Maturation Media: BrainPhys Neuronal Medium (StemCell Technologies # 05790), 1x B27 Plus Supplement (ThermoFisher Scientific # A3582801), 10 ng/ml BDNF (PeproTech # 450-02), 10 ng/ml GDNF (PeproTech # 450-10) 10 ng/ml NT-3 (PeproTech # 450-03), 1 µg/ml mouse laminin (Sigma # L2020-1MG), and 2 µg/ml doxycycline (Clontech # 631311). Half media changes were done every 3-4 days until collection at day 17.

***sgRNA cloning***

sgRNAs targeting TARDBP or a non-targeting control sgRNA were cloned into the CROP-seq vector (Addgene # 127965) and driven by the mouse U6 promoter. This vector contains EF1a promoter driving puromycin resistance and BFP expression. The CROP-seq vector was cut via restriction digest using BstXI and BlpI, and a double stranded DNA oligo with appropriate sticky end overhangs were ligated into the cut vector using DNA Ligation Kit, Mighty Mix (Takara #

6023). To generate DNA oligos with appropriate overhangs, "top" and "bottom" ssDNA oligos containing the sgRNA sequence and compatible overhangs were annealed (see sequences below):

| | sgRNA sequence | Top oligo (5'->3') | Bottom oligo (5'->3') |
|---|---|---|---|
| TARDBP sgRNA1 | GGGAAGTCAGC CGTGAGACC | TTGGGGAAGTCAGCCGTGAG ACCGTTTAAGAGC | TTAGCTCTTAAACGGTC TCACGGCTGACTTCCCC AACAAG |
| Non-targeting sgRNA (NT_02194) | GGCCGTGGGCA ACACTGTAT | TTGGGCCGTGGGCAACACTGT ATGTTTAAGAGC | TTAGCTCTTAAACATAC AGTGTTGCCCACGGCCC AACAAG |
| Non-targeting sgRNA (NT_00976) | GAAGTTACTCT ACAAAACAG | TTGGAAGTTACTCTACAAAAC AGGTTTAAGAGC | TTAGCTCTTAAACCTGT TTTGTAGAGTAACTTCC AACAAG |
| Non-targeting sgRNA (NT_02724) | GAATATGTGCG TGCATGAAG | TTGGAATATGTGCGTGCATGA AGGTTTAAGAGC | TTAGCTCTTAAACCTTC ATGCACGCACATATTCC AACAAG |

***Lentivirus generation***

Lentivirus was prepared by transfection of early passage Lenti-X 293T cells (Takara # 632180) with the sgRNA plasmid using lipofectamine 3000 using manufacturer's recommended protocol. LentiX cells were cultured in DMEM, high glucose, GlutaMAX™ Supplement (Thermo Fisher Scientific # 10566024) supplemented with 10% FBS (Thermo Fisher Scientific # A3160402). 24 hours after transfection, cells were washed with PBS and replaced with fresh media containing 1x ViralBoost (ALSTEM # VB100). Media was collected 96 hours after transfection and lentivirus was concentrated using Lenti-X concentrator (Takara Bio # 631231) using manufacturer's protocol. Lentivirus was resuspended in PBS in 1/10 of the original cell culture dish volume, aliquoted, and stored at -80°C until use.

**RNA sequencing, differential gene expression, and splicing analysis**

On day 17 of differentiation, cells were washed with PBS and lysed directly in cell culture plates with 300ul of Tri-reagent (Zymo Research # R2050-1-200). RNA was extracted from day 17 neurons using Direct-zol RNA Miniprep Kit (Zymo Research # R2051) following manufacturer's protocol, including optional DNase treatment step. RNA was quantified using a Qubit Fluorometer. Libraries were prepared using NEBNext Ultra II Directional RNA Library Prep Kit for Illumina (NEB # E7760L) with the NEBNext Poly(A) mRNA Magnetic Isolation Module (NEB # E7490L). Libraries were indexed, pooled, and sequenced (2 x 150bp) on a Novaseq 6000 using v1.5 reagents.

Paired-end fastq.gz files were trimmed for sequencing adapters using cutadapt v2.5 *(38)* and quality checked using FastQC v0.11.6*(39)*, followed by splice aware alignment against GRCh38.p13.genome.fa using STAR v2.7.3a *(40)*. Differential expression analysis was done using StringTie2 *(47)* and DESeq2 *(41)*. We ran MAJIQ (v2.1) *(42)* on aligned BAM files to perform differential splicing analysis. We set a threshold of 10% difference in percent spliced in ($\Delta\Psi$) for calling a significant change between groups. We then used custom R scripts to parse the output of the deltaPSI module to obtain the PSI ($\Psi$) and the probability of change for each junction. We defined cryptic splicing as junctions with PSI less than 5% in control samples, a $\Delta\Psi$ of more than 10%. Junctions were annotated according to the gene annotations from gencode.v37.annotation.gtf. Our splicing pipeline is implemented using Snakemake version 5.5.4 *(57)* (https://github.com/frattalab/splicing). Finally, we developed a novel pipeline to visualize and automatically categorize each mis-spliced junction from MAJIQ as cryptic exon, exon skip, likely intron retention, or canonical junction (https://github.com/NIH-CARD/proteogenomic-pipeline).

**Informatic pipeline for splicing visualization and categorization**

Given a 6 columns .csv file consisting of splice events (col1:chr#, col2:start, col3:end, col4:strand, col5:gene_name and col6:gene_id), we built a pipeline to automatically categorize each splice event as cryptic exon, exon skip, likely intron retention, or canonical junction, and to provide visual representations of each mis-splicing event in the form of sashimi plots.

***List of Transcript for probable spilling events and sashimi plots for all splicing events***

The first step in deciding the type of a splicing event is the identification of the transcript this event arises from. Using a 3-layered approach for identifying biologically relevant transcript for each splicing event, 1) Principal Isoforms from BAM file of all KD samples (using StringTie2), 2) list of APPRIS principal isoforms 1 and 3) Transcript (from EnsDB V103) with maximum number of exons and longest transcript in terms of its size (number of bp's). Using this approach, transcripts for all splicing events are identified and sashimi plots for all splicing events are generated.

***Identification of splicing event types***

Based on the overlap (using bedtools*(58)*) of the genomic coordinates of each event with the genomic coordinates of the exons of the principal Tx, an event is declared as exon skip (complete overlap between exonic boundaries), cryptic exon, intron retention, or canonical junction.

***Exon skip***

Events whose genomic coordinates are completely overlapped with two non-consecutive exons of the principal Tx are declared as exon skip events. The pipeline also differentiates between simple exon-skip events (where the event's genomic coordinates overlap with non-consecutive exons, such as 1 and 3 (net exon number difference of 2)), and multi-exon-skip events (where net exon number difference is > 2). Below is an example sashimi plot for an exon-skip event. Read coverage for each sample is shown as layered graph (different shades) for control (top panel) and KD samples (middle panel). Bottom panel (transcript lane) shows the coordinates identified by the proteogenomic pipeline, principal transcript selected and the coordinates of the mis-splicing event.

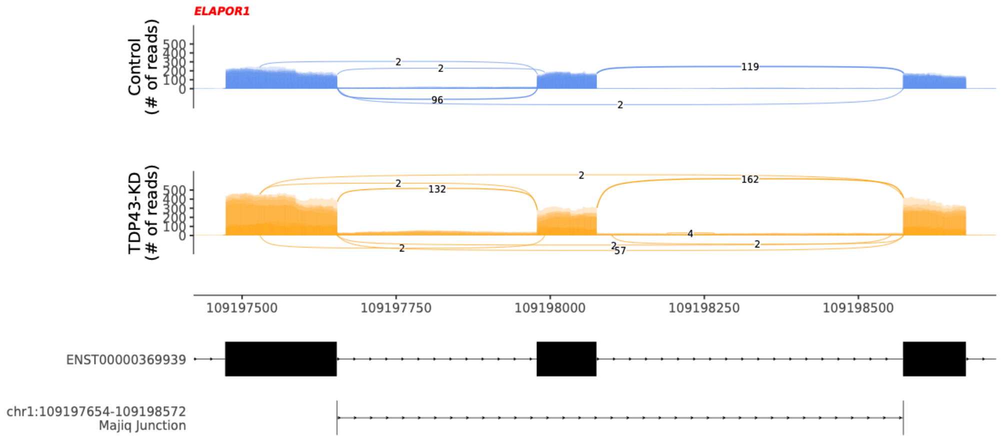


##### *Cryptic exon*

Events for which one of the genomic coordinates overlap with one of the exons in the principal Tx while the other lies in the intronic region are categorized as cryptic events. For these events, we identify genomic coordinates of the upstream and downstream exon and involved intronic region. Below is a sashimi plot for a cassette cryptic exon.

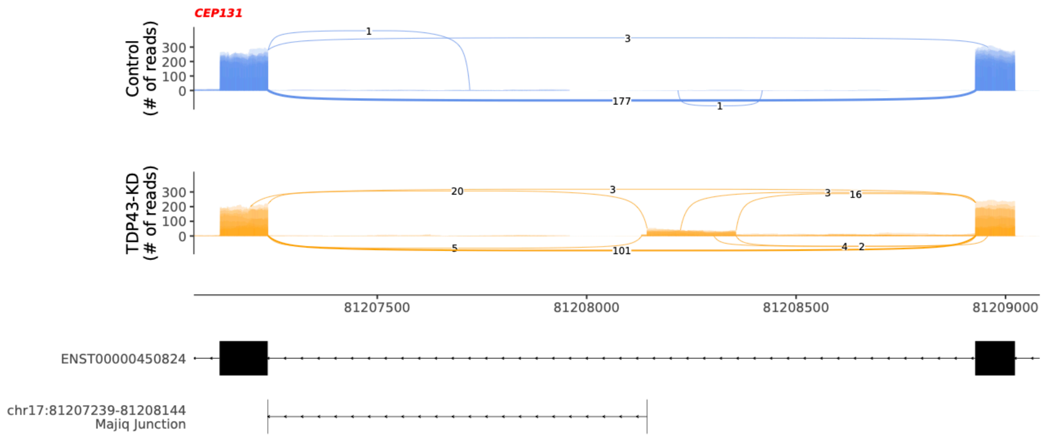


***Bed files for TDP-43 KD samples***

In order to classify whether a cryptic event is ce_inclusion, ce_extension, or IR, bed files for all BAM files for all KD samples are created. These bed files are needed to calculate coverage bed files for each probable cryptic exon.

***Coverage bed files for TDP-43 KD samples***

For each probably cryptic exon, a bed file containing the average number of reads from all KD samples for the entire intron range of interest is calculated.

***Sashimi Plots***

In order to identify genomic coordinates of a probable cryptic event, its coverage bed file is scanned from the intronic genomic coordinate of the LSV to the opposite exon. If the read coverage drops to 40% along the way, this event is classified as a CE_inclusion (cassette cryptic exon). If the coverage never drops to zero, this event is classified as CE_extension. If the coverage never drops to zero across the entire intron, then this event is classified as a probable intron retention (IR) event. Category-wise sashimi plots are also generated for user-verification.

**Long-read sequencing and data processing**

Prior library preparation, RNA integrity was assessed using Qubit$^{TM}$ RNA IQ assay kit (ThermoFisher, Q33221). 50 ng total RNAs were used to prepare cDNA-PCR libraries following manufacturer's protocol (SQK-PCS109, Oxford Nanopore Technologies (ONT)).

Sequence-specific cDNA-PCR sequencing was adapted from the ONT protocol (ss-cdna-pcr-sequencing_sqk-pcs109-CPSS_9087_v109_revO_14Aug2019-promethion). Briefly, 500 ng total RNAs were reverse-transcribed at 50°C using the Maxima H Minus Reverse Transcriptase (ThermoFisher Scientific, EP0751) in the presence of 2 pmoles of each custom sequence-specific primer (Supplementary Table S2C). The corresponding cDNA was purified with 0.5 volume of Agencourt RNAClean XP beads (Beckman, A63987) and washed with 100 µl Small Fragment Buffer (SFB). cDNA was eluted in 12 µl $H_2O$ and amplified by PCR for 15 cycles (95°C 30"; 95°C 15", 62°C 15", 65°C 8'30", x15; 65°C, 6') using the LongAmp Taq Master mix (NEB, M0287). PCR-cDNA libraries were purified using 0.5 volume of Agencourt RNAClean XP beads, washed with 100 µl Small Fragment Buffer (SFB) and eluted in 12 µl of elution buffer. Final libraries were quantified using Qubit 1X dsDNA High Sensitivity (HS) assay kit (ThermoFisher Scientific, Q33231) and analyzed on a BioAnalyzer (High Sensitivity DNA kit, Agilent, 5067-4626). Libraries were loaded onto R9.4.1 flow cells (ONT, FLO-PRO002) and ran on a Promethion device (ONT, PRO-SEQ024).

Nanopore direct RNA sequencing data were basecalled using Guppy (v3.4.5). Reads were first aligned against ribosomal sequences obtained from SILVA*(59)*. Non-ribosomal reads were subsequently mapped to the human genome (hg38) using minimap2 (version 2.17) *(43)* with parameters -a -x splice -k 12 -u b --secondary=no. Basecalled reads were also separately aligned against the human transcriptome (Ensembl version 92) using -a -x map-ont -k 12 -u f --secondary=no. Transcript identification and quantification was performed with Bambu at maximum sensitivity with parameters NDR=1, min.txScore.singleExon = 0, remove.subsetTx = TRUE, min.readCount = 2, min.readFractionByGene = 0, min.txScore.multiExon = 0, min.exonDistance = 0, min.primarySecondaryDist = 0, txScoreBaseline = 0. Only reads overlapping genes with splicing changes identified from Illumina sequencing were used.

**Ribosome profiling**

Previously published ribosome profiling libraries *(13)* were resequenced at higher depth following depletion using a Cas9-based protocol*(44)*. Reads were processed as previously described *(13)*. For analysis of periodicity (Figure 1C), reads were first processed using our standard pipeline (removing common ncRNA contaminants and PCR duplicates, and ensuring that all RFPs aligned uniquely to the genome), then FASTQs containing only uniquely aligned reads were extracted from the deduplicated genomic BAM files. The extracted FASTQs were then re-aligned to a fasta file containing just the coding sequences for the list of transcripts encoding CEs that were predicted to be translated using; the first base of each RFP was ignored (as this is often mismatched due to non-templated nucleotide addition during reverse transcription) and the rest of the footprint was required to match perfectly, through the use of harsh mismatch penalties (--norc --no-unal --very-sensitive -N 1 --mp 10000 --np 10000 --rdg 10000 --rfg 10000 -a) with Bowtie2*(60)*. The length and sub-codon position of reads aligning to CEs was then calculated. Equivalent analysis but using alignment with Bowtie2 to annotated transcripts was also performed, and the results were compared.

***De novo* peptide sequence prediction**

All MAJIQ derived cryptic splice junctions were annotated by Dasper *(45)* to separate junctions by whether they were novel acceptor, novel donor, or novel exon skipping events. Cryptic exons are those were either the donor, acceptor, or both ends of the junction is novel, whereas skipptic events are novel exon skips. In order to find the exonic regions created by these novel splice junctions, we used all exons defined in the Gencode v31, and all of the potential exonic regions defined by two different transcript assembly tools, Scallop *(46)* and Stringtie2 *(47)*. We filtered out false positive exonic regions by mapping the number of splice junctions that ended on both the 5' and 3' ends of the exonic region and those with less than 10 splice junctions on both ends across all TDP-KD samples were filtered out as being potential false positives. We mapped all novel exonic regions back to those which overlapped cryptic junctions. In order to build the possible open reading frames, we first determined which of the known transcripts overlapped with the cryptic events and could be used as 'backbone transcripts'. We then filtered out transcripts which were not expressed, with a liberal mean expression cut-off of 1 transcript per million across the all samples, determined by SALMON *(46)*. We then inserted the novel cryptic event into each of the expressed backbone transcripts, and used the genomic regions to extract the nucleotide sequences. For each nucleotide sequence, we translated into the amino acid sequence, and extracted all possible open reading frames (ORF) between every methionine and every stop codon.

#### Proteomics

##### *iNeuron (DDA, PRM) and CSF (DIA) Proteomics sample preparation*

For the iPSC-derived neurons grown and differentiated in 6-well plates, neurons were washed with ice-cold phosphate-buffered saline (PBS) three times before adding 150 ul of lysis buffer with high percentage of detergents, consisting of 50 mM Tris-HCl pH 8, 50 mM NaCl, 1% v/v SDS, 1% v/v Triton X-100, 1% v/v NP-40, 1% v/v Tween 20, 1% v/v glycerol, 1% sodium deoxycholate (w/v), 5 mM EDTA pH 8, 5 mM dithiothreitol, 5 KU benzonase and protease inhibitors. Cell lysates were collected and incubated at 65°C for 30 min, followed by addition of 10mM iodoacetamide and 30 min incubation in the dark. The automated protein digestion was performed as previously described *(48)*. Briefly, the protein input of each sample was normalized to 20 ug and was captured by the hydrophilic beads on an automated KingFisher APEX robot. The tryptic digestion was conducted at 37°C for 16 hrs on a thermomixer, and resulting peptides were dried and reconstituted in 2% acetonitrile with 0.5% trifluoroacetic acid.

For the human cerebrospinal fluid (CSF), we depleted top 14 most abundant proteins in 100 ul CSF of each patient using a rapid 10min incubation with High-Select depletion resin (Thermo Scientific). The proteins in depleted CSF were extracted and denatured by 4X lysis buffer used for iPSC-neuron lysate. The same protocol was followed as stated above. The final concentration of tryptic peptides from iPSC-neurons and CSF was normalized to 0.2 ug/ul using peptide measurement on a NanoDrop. We injected 5 ul of total peptides for proteomics analysis.

***iNeuron (DDA, PRM) and CSF (DIA) Proteomics liquid chromatography and mass spectrometry analyses***

We used three data acquisition approaches, data dependent acquisition (DDA), data independent acquisition (DIA), and parallel reaction monitoring (PRM) for liquid chromatography and tandem mass spectrometry (LC-MS/MS) analyses. The peptides were separated on an Ultimate 3000 nano-LC system coupled with a 50cm column (75 µm I.D., 2 µm $C_{18}$ particle) and were injected to a high-resolution Orbitrap Eclipse MS (Thermo Scientific). For the DDA, a 120 min effective LC gradient 2-35% liquid phase B was used. Liquid phase A was 5% DMSO in 0.1% formic acid (FA) water, and liquid phase B was 5% DMSO in 0.1% FA acetonitrile. The full

(MS1) scan was conducted 375-1400 m/z at 120k-resolution. The dynamic exclusion was set to 45 s excluding the ion after 1 time detection. The fragment (MS2) scan was set to cycle time 3 s, isolation window 1.6 m/z, HCD collision energy 30%, and the fragments were detected in linear ion trap at rapid scan rate with auto maximum injection time. For the DIA, a 90 min effective LC gradient 2-35% liquid phase B was used. The MS1 scan was conducted 390-1010 m/z at 120k-resolution. For MS2 scan, we used an isolation window at 8 m/z in a range of 400-1000 m/z with 30% HCD collision energy. The fragments were detected in the orbitrap with a 30k resolution and 3 s loop control. For the PRM, a 60 min effective LC gradient 2-35% liquid phase B was used. The MS1 scan was conducted 350-2000 m/z at 120k-resolution. The MS2 was conducted based on mass list table generated by Skyline using orbitrap with a 30k resolution and 3 s loop control.

***iNeuron (DDA, PRM) and CSF (DIA) Proteomics database search and statistical analyses***

For DDA-based discovery proteomics, we used PEAKS studio (v10.6) with a customized database containing uniport-human proteome reference (UP000005640) and open reading frames (ORFs*) in silico* translated from mis-spliced junctions identified by total RNA sequencing. Trypsin was chosen as digestion enzyme (allowing semi-tryptic digestion), and mass error tolerance was set to 15ppm, and 3 mis-cleavages was allowed. For DIA database search, direct DIA-library was used in Spectronaut (v16) *(51)* with the same customized FASTA database, and searching parameters were same as DDA. For the targeted proteomics, MS raw files of PRM or DIA were directly loaded to Skyline (daily version) *(50)* for quantification and visualization. Specifically, only common y-ions identified in both light and heavy peptides were used for quantification and dot product estimation. The light peptide intensity was normalized by its heavy labeled counterpart. The statistical analysis was conducted in R studio (v4.3), the simple two-side t-test was used for comparison, and p-values were adjusted for multiple-comparisons using Benjamini-Hochberg procedure. The differentiation expression analysis was carried out in Spectronaut (v16) *(51)* where log fold changes and q-value were generated for volcano plot analysis.

***iNeuron (PRM) and CSF (DIA) Proteomics addition of heavy isotopic labeled peptides***

The standard peptides were 8-15mers (i.e., amino acid residues in length) identified in discovery proteomics or predicted by PeptideRank (http://wlab.ethz.ch/peptiderank/) where trypsin was chosen as the protease. To make their heavy isotopic labeled counterparts, we labeled their lysine (i.e., $^{13}C_6{}^{13}N_2$, Lys8) or arginine (i.e., $^{13}C_6{}^{13}N_4$, Arg10) at c-terminus except the semi-tryptic peptides which were labeled at the closest lysine or arginine to c-terminus. The heavy labeled peptides were reconstituted to 0.05 to 1pmol/ul based on their signal abundance in mass spectrometry comparing with their light counterparts (endogenous peptides) and were pooled. The iRT standards (Biognosys) and pooled heavy standards were spiked into the digested peptides at 1:10 ratio for targeted proteomics.

***iNeuron DIA Proteomics sample preparation***

Cell pellets of five control and five TDP-43 knockdown (KD) samples were collected in 2ml Eppendorf tubes and snap-frozen in liquid nitrogen. On preparation for MS, samples were thawed on ice before adding 200 µl lysis buffer (2% SDS, 100 mM HEPES, pH8, 50 mM DTT). Samples were then vortexed (five times) prior to sonication (Bioruptor Plus) for 10 cycles (30 s ON/60 s OFF) at high setting, at 20°C. Reduction (15 min, 45°C) was followed by alkylation with 20 mM iodoacetamide (IAA) for 30 min at room temperature in the dark. Protein amounts were quantified by EZQ™ (Invitrogen) and 200µg of protein from each sample were taken along for digestion. Proteins were precipitated overnight at −20°C after addition of a 4× volume of ice-cold acetone. The following day, the samples were centrifuged at 20,800 *g* for 30 min at 4°C and the supernatant carefully removed. Pellets were washed twice with 1 ml ice-cold 80% (v/v) acetone in water then centrifuged at 20,800 *g* at 4°C. They were then allowed to air-dry before addition of 100 µl of digestion buffer (1M Guanidine, 100 mM HEPES, 100mM HEPES, pH8), followed by sonication (as above) and addition of LysC (Wako) at 1:100 (w/w) enzyme:protein for 4 h at 37°C with shaking (Eppendorf ThermoMixer®C, thermoblock for 1.5 ml tubes, at 1,000 rpm for 1 h, then 650 rpm). Samples were then diluted 1:1 with Milli-Q water, and trypsin (Promega) added at the same enzyme to protein ratio. Samples were further digested overnight at 37°C with shaking (650 rpm). The following day, digests were acidified by the addition of TFA to a final concentration of 2% (v/v) and then desalted with Waters Oasis® HLB µElution Plate 30 µm (Waters Corporation, Milford, MA, USA) in the presence of a slow vacuum. In this process, the columns were conditioned with 3 × 100 µl solvent B (80% (v/v) acetonitrile; 0.05% (v/v) formic acid) and equilibrated with 3 × 100 µl solvent A (0.05% (v/v) formic acid in Milli-Q water). The samples were loaded, washed 3 times with 100 µl solvent A, and then eluted into 0.2-ml PCR tubes with 50 µl solvent B. The eluates were dried down with the speed vacuum centrifuge and dissolved at a concentration of 1 µg/µl in reconstitution buffer (5% (v/v) acetonitrile, 0.1% (v/v) formic acid in Milli-Q water). Samples were spiked with iRT peptides (Biognosys, Switzerland) and analyzed as described below.

***iNeuron DIA Proteomics data processing***

Direct DIA search was performed against a human FASTA database (SwissProt, release 2016_01) *(61)* and a list of common contaminants using Pulsar engine in Spectronaut Professional+ (version

14.9, Biognosys AG, Schlieren, Switzerland) *(51)*. The following modifications were included in the search: Carbamidomethyl (C) (Fixed) and Oxidation (M)/Acetyl (Protein N-term; Variable). A maximum of 2 missed cleavages for trypsin were allowed. The identifications were filtered to satisfy FDR of 1% on peptide and protein levels. Precursor matching, protein inference, and quantification were performed in Spectronaut using default settings. The contrast table (differential expression analysis) was then exported to perform further data visualization (i.e., volcano plot) using R/Bioconductor.

***iNeuron DIA Proteomics mass spectrometry analysis***

Peptides were separated in trap/elute mode using the nanoAcquity MClass Ultra-High Performance Liquid Chromatography system (Waters, Waters Corporation, Milford, MA, USA) equipped with a trapping (nanoAcquity Symmetry C18, 5 µm, 180 µm × 20 mm) and an analytical column (nanoAcquity BEH $C_{18}$, 1.7 µm, 75 µm × 250 mm). Solvent A was water and 0.1% formic acid, solvent B was acetonitrile, and 0.1% formic acid. 1 µl of the sample (∼1 µg on column) was loaded with a constant flow of solvent A at 5 µl/min onto the trapping column. The trapping time was 6 min. Peptides were eluted via the analytical column with a constant 0.3 µl/min flow. During the elution, the percentage of solvent B increased in a nonlinear fashion from 0–40% in 120 min. The total run time was 145 min, including equilibration and conditioning. The LC was coupled to an Orbitrap Exploris 480 (Thermo Fisher Scientific, Bremen, Germany) using the Proxeon nanospray source. The peptides were introduced into the mass spectrometer via a Pico-Tip Emitter 360-µm outer diameter × 20-µm inner diameter, 10-µm tip (New Objective) heated at 300 °C, and a spray voltage of 2.2 kV was applied. The capillary temperature was set at 300°C. The radiofrequency ion funnel was set to 30%. For DIA data acquisition, full-scan mass spectrometry (MS) spectra with a mass range of 350–1650 m/z were acquired in profile mode in the Orbitrap with a resolution of 120,000 FWHM. The default charge state was set to 3+. The filling time was set at a maximum of 60 ms with a limitation of $3 \times 10^6$ ions. DIA scans were acquired with 40 mass window segments of different widths across the MS1 mass range. Higher collisional dissociation fragmentation (stepped normalized collision energy; 25, 27.5, and 30%) was applied, and MS/MS spectra were acquired with a resolution of 30,000 FWHM with a fixed first mass of 200 m/z after accumulation of $3 \times 10^6$ ions or after filling time of 35 ms (whichever occurred first). Data were acquired in profile mode. For data acquisition and processing of the raw data Xcalibur 4.3 (Thermo) and Tune version 2.0 were used.

***CSF TmT Proteomics sample preparation***

Briefly, 150 µl of CSF from each of the 24 individual CSF samples was incubated with 200 µl of High Select Top14 Abundant Protein Depletion Resin (Thermo Fisher Scientific, A36372) at room temperature in centrifuge columns (Thermo Fisher Scientific, A89868) essentially as previously described *(56)*. After 15 min of rotation, the samples were centrifuged at 1000*g* for 2 min. Sample flow-throughs were reduced and alkylated with 5 µl of 0.5 M tris-2(-carboxyethyl)-phosphine and 25 µl of 0.4 M chloroacetamide at 90°C for 10 min, followed by water bath sonication for 15 min. The samples were dried by speed vacuum (Labconco), then resolved in 150ul of 6M urea buffer [6 M urea and 75 mM sodium phosphate (pH 8.5)]. Protein concentration was assessed by bicinchoninic acid (BCA) method according to the manufacturer's protocol (Thermo Fisher Scientific). Immunodepleted CSF (125 µl) from all samples was digested with lysyl endopeptidase (LysC) and trypsin. Briefly, LysC (5 µg; Wako) was used for overnight digestion at room temperature. Samples were then diluted to 1 M urea with 50 mM ammonium bicarbonate (ABC). An equal amount (5 µg) of trypsin (Promega) was added, and the samples were subsequently incubated for 12 hours. The digested peptide solutions were acidified to a final concentration of

1% formic acid (FA) and 0.1% trifluoroacetic acid (TFA), followed by desalting with 30 mg of HLB C18 columns (Waters) as described previously *(56)*. The peptides were subsequently eluted in 1 ml of 50% acetonitrile (ACN). To normalize protein quantification across batches, 150 μl of aliquots from all 24 CSF samples were combined to generate a pooled sample, which was then aliquoted into 3 global internal standard (GIS) with 850 μl each. All individual samples and the pooled standards were dried by speed vacuum (Labconco).

All 24 samples and 3 GIS samples were divided into three batches, labeled using an 11-plex TMT kit (Thermo Fisher Scientific, lot no.RF234620), and derivatized as previously described *(56)*. See the Supplementary Table S6A for sample to batch arrangement. Nine of the 11 TMT channels were used for labeling: 127N, 128N, 128C, 129N, 129C, 130N, 130C, 131N, and 131C. Briefly, 5 mg of each TMT reagent was dissolved in 256 μl of anhydrous ACN. Each CSF peptide digest was resuspended in 50 μl of 100 mM triethylammonium bicarbonate (TEAB) buffer, and 20.5 μl of TMT reagent solution was subsequently added. After 1 hour, the reaction was quenched with 4 μl of 5% hydroxylamine (Thermo Fisher Scientific, 90115) for 15 min. After labeling, the peptide solutions were combined according to the batch arrangement. Each TMT batch was desalted with 100 mg of Sep-Pak C18 columns (Waters) and dried by speed vacuum (Labconco).

To enhance the depth of the discovery CSF proteome, these TMT labeled samples were subjected to high-pH fractionation as previously described *(56)*. TMT-labeled peptides (~220 μg) from each discovery sample were dissolved in 100 μl of loading buffer [1 mM ammonium formate in 2% (v/v) ACN], injected completely with an autosampler, and fractionated using a ZORBAX 300Extend-C18 column (2.1 mm by 150 mm, 3.5 μm; Agilent Technologies) on an Agilent 1100 HPLC (high-performance liquid chromatography) system monitored at 280 nm. A total of 96 fractions were collected over a 60-min gradient of 100% mobile phase A [4.5 mM ammonium formate (pH 10) in 2% (v/v) ACN] from 0 to 2 min, 0 to 12% mobile phase B [4.5 mM ammonium formate (pH 10) in 90% (v/v) ACN] from 2 to 8 min, 12 to 40% mobile phase B from 8 to 36 min, 40 to 44% mobile phase B from 36 to 40 min, 44 to 60% mobile phase B from 40 to 45 min, and 60% mobile phase B until completion with a flow rate of 0.4 ml/min. The 96 fractions were collected with an even time distribution and pooled into 24 fractions. All fractions were dried down by vacuum centrifugation in a speedvac.

***CSF TmT Proteomics mass spectrometry analysis***

Each fraction was brought up in 10 ul of loading buffer (0.1% FA and 0.03 % TFA) and 2 ul was loaded onto an in-house packed analytical column (75um x 50 cm packed with 1.9um Dr. Maisch ReproSil-Pur C18 beads). Elution was carried out across 100 mins at a flow rate of 250nl/min using an EasyNlc 1200 (ThermoFisher). The gradient went from 1% B (0.1% FA in ACN) to 40% B. Mass spectra were collected on a Orbitrap Fusion Lumos mass spectrometer. The mass spectrometer was set to collect at top speed with a cycle time of 3 seconds. Full scan were collected with a range of 375-1500 m/z, 120k resolution, an AGC of 40000 and a maximum injection time of 50ms. Tandem scans were collected at 50k resolution with an isolation window of 1.2 m/z, collision energy of 36%, an AGC target of 50k and a max injection time of 86ms.

***CSF 2-D Proteomics sample preparation***

A pooled sample was made by mixing 28 ALS csf samples with equal volume, whose trait was attached. The pool sample(800ul) was reduced and alkylated with 16 µl of 0.5 M tris-2(-carboxyethyl)-phosphine and 80 µl of 0.4 M chloroacetamide at 90°C for 10 min, followed by water bath sonication for 10 min. sample was diluted with 896 µl of 8 M urea buffer [8 M urea and 100 mM NaHPO4 (pH 8.5)] to a final concentration of 4 M urea. LysC (300 mAU, ) was used for overnight digestion at room temperature. 120 µg of trypsin (Promega) was added after the sample was diluted to 1 M urea with 50 mM ammonium bicarbonate (ABC). The digestion was carried out for another 12 hours at room temperature. The digested peptide solution was acidified to a final concentration of 1% formic acid (FA) and 0.1% trifluoroacetic acid (TFA) (66), followed by desalting with 2 of 30 mg of HLB C18 columns (Waters) as described previously (cite ‘ Large-scale Proteomic Analysis of Alzheimer’s Disease Brain and Cerebrospinal Fluid Reveals Early Changes in Energy Metabolism Associated with Microglia and Astrocyte Activation’). The peptides were subsequently eluted in 2 ml of 50% acetonitrile (ACN) then dried by speed vacuum (Labconco).

***CSF 2-D Proteomics high-pH fractionation***

Dried samples were re-suspended in high pH loading buffer (0.07% vol/vol $NH_4OH$, 0.045% vol/vol FA, 2% vol/vol ACN) and loaded onto a Water’s BEH (2.1mm x 150 mm with 1.7 µm beads). An Thermo Vanquish UPLC system was used to carry out the fractionation. Solvent A consisted of 0.0175% (vol/vol) $NH_4OH$, 0.01125% (vol/vol) FA, and 2% (vol/vol) ACN; solvent B consisted of 0.0175% (vol/vol) $NH_4OH$, 0.01125% (vol/vol) FA, and 90% (vol/vol) ACN. The sample elution was performed over a 25 min gradient with a flow rate of 0.6 mL/min with a gradient from 0 to 50% B. A total of 192 individual fractions were collected, consolidated down to 96 fractions and dried by speed vacuum (Labconco).

***CSF 2-D Proteomics mass spectrometry analysis***

Each of the 96 high-pH peptide fractions was resuspended in loading buffer (15ul of 0.1% FA, 0.03% TFA, 1% ACN), and 3ul were separated on a Water’s CSH C18 column (1.7um C18 150um x 15 cm) by an Easy-nLC 1200 system (Thermo Fisher Scientific) and monitored on an Orbitrap Fusion mass spectrometer (Thermo Fisher Scientific). Elution was performed over a 45 min gradient at a rate of 1250 nl/min with Buffer B ranging from 1% to 99% (Buffer A: 0.1% FA in water, Buffer B: 80% ACN in water and 0.1% FA). The mass spectrometer was set to acquire data in top speed mode with 3 s cycles. Full MS scans were collected at a resolution of 120,000 (300-1500 m/z range, 4 × 10e5 AGC, 50 ms maximum ion time). All HCD MS/MS spectra were acquired at a resolution of 15,000 (0.7 m/z isolation width, 35% collision energy, 54 ms maximum ion time). Dynamic exclusion was set to exclude previously sequenced peaks for 20 s within a 10 ppm isolation window. Only charge states from 2+ to 5+ were chosen for tandem MS/MS.

**Splicing analysis of postmortem brain tissue**

FACS-sorted frontal cortex neuronal nuclei were obtained from the Gene Expression Ombius (GEO) GSE126543 and aligned, as per *(13)*. Briefly, samples were quality trimmed using Fastp aligned to the GRCh38 genome using STAR (v2.7.0f) *(40)* with gene models from GENCODE v31. Our alignment pipeline is implemented in Snakemake version 5.5.443 and available at: https://github.com/frattalab/rna_seq_snakemake. STAR's splice junction output tables were then clustered and converted into PSI metrics using Dasper *(45)*. As splicing tools can be prone to one-off errors for exact splice junction coordinates, the 340 *bona fide* splicing events from MAJIQ were manually curated against the splice junctions output by STAR to confirm the absence or presence of the event in the FACs-sorted nuclei.

Our analysis of postmortem brain tissue from NYGC contains 472 neurological tissue samples from 286 individuals samples from the NYGC ALS dataset, including non-neurological disease controls, FTLD, ALS, FTD with ALS (ALS-FTLD), or ALS with suspected Alzheimer's disease (ALS-AD). Patients with FTD were classified according to a pathologist's diagnosis of FTD with

TDP-43 inclusions (FTLD-TDP), or those with FUS or Tau aggregates. ALS samples were divided into the following subcategories using the available Consortium metadata: ALS with or without reported SOD1 or FUS mutations. ALS-TDP was categorized as all non-SOD1 or FUS ALS samples, but postmortem TDP-43 inclusions pathology was not systematically measured in ALS samples. Sample processing, library preparation, and RNA-seq quality control are described previously *(10, 62)*. RNA was extracted from flash-frozen postmortem tissue using TRIzol (Thermo Fisher Scientific) chloroform and 500 ng of the total RNA was used to create libraries for RNA-Seq with the KAPA Stranded RNA-Seq Kit with RiboErase (KAPA Biosystems) for ribosomal RNA depletion. The libraries had an average insert size of 375 bp and y were sequenced either on an Illumina HiSeq 2500 (125 bp paired end) or an Illumina NovaSeq (100 bp paired end). The samples had a median sequencing depth of 42 million read pairs with a range of 16 to 167 million read pairs.

All samples underwent the same processing, which included adapter trimming with Trimmomatic and alignment to the GRCh38 genome with STAR (2.7.2a) *(40)* with indexes from GENCODE v30. Quality control was conducted thoroughly to validate sex and tissue type of samplew using SAMtools *(63)* and Picard Tools (Broad Institute, GitHub Repository: https://broadinstitute.github.io/picard/. A modified form of our splice junction parsing pipeline *(13)* was used to find read counts in the NYGC dataset for the 340 *bona fide* splicing events from MAJIQ. We extracted all junctions from the NYGC samples which overlapped the 340 *bona fide* splice events and manually curated them to ensure that no splice junctions were missed due to one-off errors on exact splice coordinates. For each splice junction, to calculate the area under the curve (AUC) for classification performance between TDP-43-proteinopathy and non-TDP-43 proteinopathy samples, the read count for each splice junction was first library-sized normalized and then converted to z-scores across samples. Meta-scores were created as sum of the z-scores across either all cryptic junctions, only those junctions which have a positive predictive value above 0.6, or predictive junctions excluding *STMN2*. AUC scores were calculated in R version 4.2.1, using the *pROC* package *(52)*.

**qRT-PCR validation of cryptic exons**

Frontal cortex tissues from individuals with neuropathologically confirmed FTLD-TDP and those without neuropathological features were provided by the Mayo Clinic Florida Brain Bank. Written informed consent was provided by all participants or their family members, and all protocols were approved by the Mayo Clinic Institution Review Board and Ethics Committee. Sample size was determined based on the availability of tissue in our brain bank. Levels of the cryptic transcript variants were determined using complementary DNA (cDNA) resulting from 500 ng of RNA (RNA integrity, RIN $\geq$ 7.0) that was available from a previous study *(64)*. CRISPRi-i3Neuron iPSCs (i3877 N) cDNA was generated from our previous publication *(10)*, in which TDP-43 is downregulated to about 50%. Quantitative real-time PCRs (qRT-PCR) were conducted using SYBR GreenER qPCR SuperMix (Invitrogen) for all samples in triplicates. qRT-PCR were run in a QuantStudio™ 7 Flex Real-Time PCR System (Applied Biosystems). Relative quantification was determined using the $\Delta\Delta$Ct method and normalized to the endogenous controls *GAPDH* and *RPLP0*. Primer efficiency was verified for each cryptic exon variants prior running the qRT-PCRs. See Supplementary Table S3E for primers. To compare *tSTMN2* RNA between controls and FTLD-TDP cases, unpaired Mann-Whitney tests were performed. To compare *tSTMN2* RNA between i3Neurons treated with control sgRNA (sgControl) and the two different sgRNAs against *TARDBP* (sgTARDBP-1 and siTARDBP-8), we used one-way ANOVA. All statistical analyses were done using GraphPad Prism 9 (GraphPad Software). For each figure the type of analysis used, and the number of subjects is indicated in the figure and/or legend.

**MYO18A-CE western blot**

iNeuron pellets were lysed in Co-IP buffer (50 mM Tris–HCl, pH 7.4, 300 mM NaCl, 1% Triton X-100, 5 mM EDTA) plus both protease and phosphatase inhibitors, sonicated on ice, and then centrifuged at 16,000 × g for 20 min. Supernatants were saved as cell lysates. The protein concentration of lysates was determined by BCA assay, and samples were then subjected to Western blot analysis. Equal amounts of protein were loaded into 10-well 3-8% Tris-Acetate (MYO18A-CE) or 4-20% Tris-Glycine gels (TDP-43 and GAPDH) (ThermoFisher). After transferring proteins to membranes, membranes were blocked with 5% nonfat dry milk in Tris-buffer saline (TBS) plus 0.1% Tween 20 (TBST) for 1 hour, then incubated with anti-rabbit MYO18A-CE antibody (1:500), anti-rabbit TDP-43 antibody (1:1000, Proteintech, 12892-1-AP) or anti-mouse GAPDH antibody (1:5000, Meridian Life Science, H86504M) overnight at 4°C. Membranes were washed in TBST and incubated with donkey anti-rabbit or anti-mouse IgG antibodies conjugated to horseradish peroxidase (1:5000; Jackson ImmunoResearch) for 1 h at room temperature. Protein expression was visualized by enhanced chemiluminescence treatment and exposure to Amersham ImageQuant 800. MYO18A-CE antibody to detect the cryptic sequence in MYO18A was generated by Labcorp by immunizing rabbits with a peptide including the complete 20 residue neoepitope (VKEEDKTLPKPGSPGKEEGA).

**HDGFL2 western blot**

iNeurons were maintained in BrainPhys media (StemCell Technlogies) supplemented with 1x B27 (Thermo Scientific), 1x N2 (Thermo Scientific), 20 ng/ml BDNF (PeproTech) and 20 ng/ml GDNF (PeproTech) for 30 days on poly-D-lysine coated 12 well plates. The cells were harvested with 4x

LDS NuPage buffer (Thermo Scientific) and protein concentration was measured with the Pierce 660nm Protein Assay Kit (Thermo Scientific). Equal amounts of protein were loaded in 7% Bis-Tris gels and run with MOPS buffer (Thermo Scientific) then transferred to 0.2 um PVDF membranes (Amersham GE Healthcare). Primary antibodies used were rabbit anti-HDGFL2 (1:1000 Sigma HPA 044208), mouse anti-TDP-43 (1:5000 Abcam ab104223) and rat anti-Tubulin (1:5000 Millipore MAB1864). Western blots were developed on a Bio-Rad ChemiDoc and quantified with ImageJ.

**Affinity purification mass spectrometry**

Each 10-cm dish of cells was washed three times with ice-cold phosphate-buffered saline (PBS; Lonza Cat. #17-516F) and lysed with 0.5 mL of ice-cold lysis buffer containing 50 mM TrisHCL, pH 8, 150 mM NaCl, 0.1% TritonX-100, and 1x protease inhibitor cocktail (Roche SKU 5892791001). Each dish was rocked for 15 min at 4°C before collecting cell lysates and centrifuging at 20,000g for 15 min at 4°C. The remaining lysate supernatants were collected. Antibodies for the appropriate control or experimental condition were added to each sample and rotated overnight at 4°C for 16 hours. Sample protein concentrations were evaluated using a detergent compatible protein assay (DCA) (Bio-Rad, Hercules, CA, Cat. #5000111) and normalized to 0.4 µg/µL. Automated affinity purification was then performed for 4 hours on a KingFisher Flex Purification System (Thermo Scientific) at 4°C. Eluates were transferred to a ThermoMixer and incubated at 37°C for 16 hours. Peptides were then dried using a SpeedVac vacuum concentrator (Thermo Scientific) and reconstituted in 2% acetonitrile and 0.4% trifluoroacetic acid (TFA). The peptide concentrations were evaluated on a Denovix DS-11 FX Spectrophotometer/Fluorometer (peptide mode, acquisition wavelength 215 nm, E 0.1% (mg/mL), correction factor of 25.99) and normalized to 0.1 µg/µL. Analysis of 5 µL of each normalized peptide sample was conducted via LC-MS/MS. Precursor matching, protein inference, and quantification were performed in Spectronaut using the DirectDIA workflow in default settings. Differential abundance analysis was carried out in the Spectronaut software 16.2, generating log fold changes and q-values by using a two-sided t-test and Benjamini-Hochberg adjustment for multiple comparisons. R software version 4.2.3 was used to data visualization.

*Please note: all supplementary files and tables can be accessed here (https://shorturl.at/HJORU)*
